# A genome engineered tool set for *Drosophila* TGF-β/BMP signaling studies

**DOI:** 10.1101/2024.07.02.601693

**Authors:** Clara-Maria Ell, Abu Safyan, Mrinal Chayengia, Manuela M. M. Kustermann, Jennifer Lorenz, Melanie Schächtle, George Pyrowolakis

## Abstract

Ligands of the TGF-β/BMP superfamily are critically involved in the regulation of growth, patterning and organogenesis and can act as long-range morphogens. Essential for understanding TGF-β/BMP signaling dynamics and regulation are tools that allow monitoring and manipulating pathway components expressed at physiological levels and endogenous spatiotemporal patterns. We used genome engineering to generate a comprehensive library of endogenously epitope-or fluorescently-tagged versions of receptors, co-receptors, transcription factors and key feedback regulators of the *Drosophila* BMP and Activin signaling pathways. We demonstrate that the generated alleles are biologically active and can be utilized for assessing tissue and subcellular distribution of the corresponding proteins. Further, we show that the genomic platforms can be used for *in locus* structure-function and *cis*-regulatory analyses. Finally, we present a complementary set of protein binder-based tools, which allow visualization as well as manipulation of the stability and subcellular localization of epitope-tagged proteins, providing new tools for the analysis of BMP signaling and beyond.

## Introduction

Transforming growth factor beta/Bone Morphogenetic Protein (TGF-β/BMP) signaling is crucial for animal development and homeostasis and is deregulated in various pathologies and diseases (Jia and Meng, 2021; Massagué and Sheppard, 2023). Despite context-dependent differences in complexity and regulation, the core pathway of the canonical TGF-β/BMP signaling that transmits the information from extracellular ligands to the nucleus of signal-receiving cells is relatively simple and evolutionary highly conserved. Ligands of the TGF-β, BMP, and Activin families assemble into dimers and bind to extracellular domains of membrane-bound type I and type II receptor serine-threonine kinases. Upon ligand binding, type II receptors phosphorylate a glycine and serine-rich juxtamembrane domain (GS-domain) of type I receptors. Activated type I receptors then phosphorylate C-terminal serine residues of receptor-associated Smads (R-Smads), which associate with common Smad (co-Smad) and accumulate in the nucleus. Here, the Smad complexes directly bind to DNA and regulate, together with other transcription factors and co-regulators, target gene transcription.

The high evolutionary conservation allowed the use of model organisms such as *Drosophila* to identify many components of TGF-β/BMP signaling and to unravel key concepts of the signal transduction and regulation of the pathway over the past years (Akiyama et al., 2024). In *Drosophila*, ligands of the BMP and Activin branches of the TGF-β/BMP superfamily are required throughout fly development and homeostasis to regulate processes such as cell proliferation, cell differentiation, cell migration and apoptosis (Upadhyay et al., 2017). The best-studied BMP ligand in *Drosophila* is Decapentaplegic (Dpp), the fly homolog of vertebrate BMP2 and BMP4. Dpp, which often forms a heterodimer with Glass bottom boat (Gbb) or Screw (Scw), the other *Drosophila* BMPs, plays essential roles in embryonic dorso-ventral axis formation, germline and intestinal stem cell maintenance, patterning and growth of larval imaginal discs and patterning of the follicular epithelium during oogenesis (Akiyama et al., 2024; Upadhyay et al., 2017). Detailed studies in some of these tissues underlined the necessity of precise spatiotemporal regulation of Dpp/BMP signaling activity (Bier and De Robertis, 2015; Montanari et al., 2022; Wilcockson et al., 2017). This is particular evident in the context of larval wing development, were Dpp, in parallel to its role in promoting growth, acts as a morphogen to provide positional information (Affolter and Basler, 2007; Ashe and Briscoe, 2006; Hamaratoglu et al., 2014; Kicheva and Briscoe, 2023; Stapornwongkul and Vincent, 2021). Decades of research in this tissue, which has widely served as a paradigm for morphogen signaling, highlighted a number of determinants and interactions shaping the BMP activity gradient. While localized expression of *dpp* in a stripe of anterior cells at the anterior-posterior compartment boundary and dispersion from this source are thought to underlie gradient formation, the exact mechanisms by which Dpp spreads into both compartments are not completely understood. Nevertheless, it has become evident that the sum of multiple interactions acting at distinct levels regulate gradient shape and transcriptional output. At the membrane surface, the distribution, levels and activity of the main receptor Thickveins (Tkv), the glypican Division abnormally delayed (Dally) and the Dally-binding protein Pentagone (Pent) are critically involved in the distribution and activity of Dpp (Akiyama et al., 2024). Within the cell’s cytoplasm, Daughters against dpp (Dad) serves as an inhibitory Smad (I-Smad) and feedback regulator of the pathway to regulate phosphorylation levels of the R-Smad Mothers against Dpp (Mad). Finally, the output of the gradient in the nucleus, i.e. the transcriptional activation of BMP target genes, is controlled by the transcriptional repressor Brinker (Brk), which counteracts Smad-dependent gene activation and is itself directly repressed by the nuclear Smad gradient (Affolter and Basler, 2007; Hamaratoglu et al., 2014). A similarly high necessity for spatiotemporal regulation of BMP signaling activity has been illustrated, or is expected, in other developmental and homeostatic contexts, including BMP-dependent embryonic axis determination, intestinal and germline stem cell maintenance and regeneration, synaptogenesis as well as follicle cell patterning (Akiyama et al., 2024; Bier and De Robertis, 2015; Hamaratoglu et al., 2014; O’Connor et al., 2006; Pyrowolakis et al., 2017; Umulis et al., 2009; Wilcockson et al., 2017).

Analyses of BMP signaling in the above-mentioned contexts require reagents and tools that allow for monitoring and manipulating the involved proteins at physiological levels and endogenous expression patterns. We present here a collection of genome engineered core components of the pathway that allows efficient epitope and fluorescent protein tagging and visualization of tissue and cellular distribution of the corresponding proteins. Furthermore, we show how the genomic platforms can be used to address isoform and structural requirements for protein function. To this end, we demonstrate that a hitherto uncharacterized Mad isoform, Mad-PB, is positively regulated by BMP signaling in the wing disc and is essential for proper adult wing size. In addition, we show that membrane anchoring, but not necessarily GPI-mediated anchoring, of the glypican Dally is required for its biological activity. Lastly, we present a complementary set of tools based on a nanobody against the HA epitope tag, which, along with previously established nanobody-based tools against GFP, can be used for trapping, mislocalizing or degrading tagged proteins of our library.

## Results

### Generation of endogenously tagged TGF-β/BMP components

We employed a two-step protocol to generate endogenously epitope or fluorescently-tagged TGF-β/BMP signaling components (Fig. 1 and Fig. S1). In the first step, we used genome engineering to replace a selected region of the target gene by a sequence containing an attP site along with a loxP–flanked selection cassette. The exact strategy and the choice of the introduced deletion were individually adapted to take into account the genomic constraints and architecture of the respective target locus as well as the domain structure of the corresponding protein (Fig. 1B and Fig. S1A). Overall, we generated chromosomal lesions in 14 genes of the *Drosophila* TGF-β/BMP signaling pathway. The introduced deletions were verified both genetically and molecularly (see Material and Methods and Fig. S1B,C) and comprise a collection of new, molecularly defined null alleles of the corresponding genes. In the second step, we used ФC31/attB integration to reinstate the genomic locus with epitope or fluorescently-tagged versions of the gene. Depending on the target gene and the introduced deletion, the strategy varied to reintegrate single exons, full-length cDNA versions of the gene, or extended genomic sequences including introns and/or flanking intergenic sequences. We introduced a number of tags including eGFP, YFP, CFP, HA (in single or triple copies), V5 and FGT (a fusion of 3xFLAG, eGFP and 2xTY1 described in (Sarov et al., 2016)). Depending on the protein, tags were introduced either at the C-terminus (receptors) or at the N-terminus (most transcription factors) (Fig. 1C). For proteins that contain signal peptides and do not tolerate C-terminal modifications (Pent and glypicans), tags were introduced internally, close to the N-terminus but downstream of the signal peptidase cleavage site. The modified genes were tested for their ability to substitute for the function of the corresponding endogenous gene. At least one (and up to five) of our epitope or fluorescently-tagged versions for each gene is functional, based on the criterion that flies carrying the introduced modifications in homozygosity are viable, fertile and do not display gross phenotypic abnormalities (Fig. 1C). All modified components have been reported to contribute to wing development, a tissue that is particular sensitive to perturbations of TGF-β/BMP signaling. To address the biological activity of the tagged proteins, we scored for morphological abnormalities in adult wings that carry the modified components in homozygosity (Fig. S2). In the majority of the cases, wings were normal in size and morphology, indicating that the modified protein is fully functional. Notable exceptions are wings of flies homozygous for Tkv-YFP, which occasionally displayed venation defects including truncations at the distal tip and local thickening of veins. Since these phenotypes were only present in Tkv-YFP but not in flies carrying the other modified versions of Tkv, we conclude that they are not caused by the tagging approach or the position of the tag but rather by the nature of the tag.

**Figure 1.**
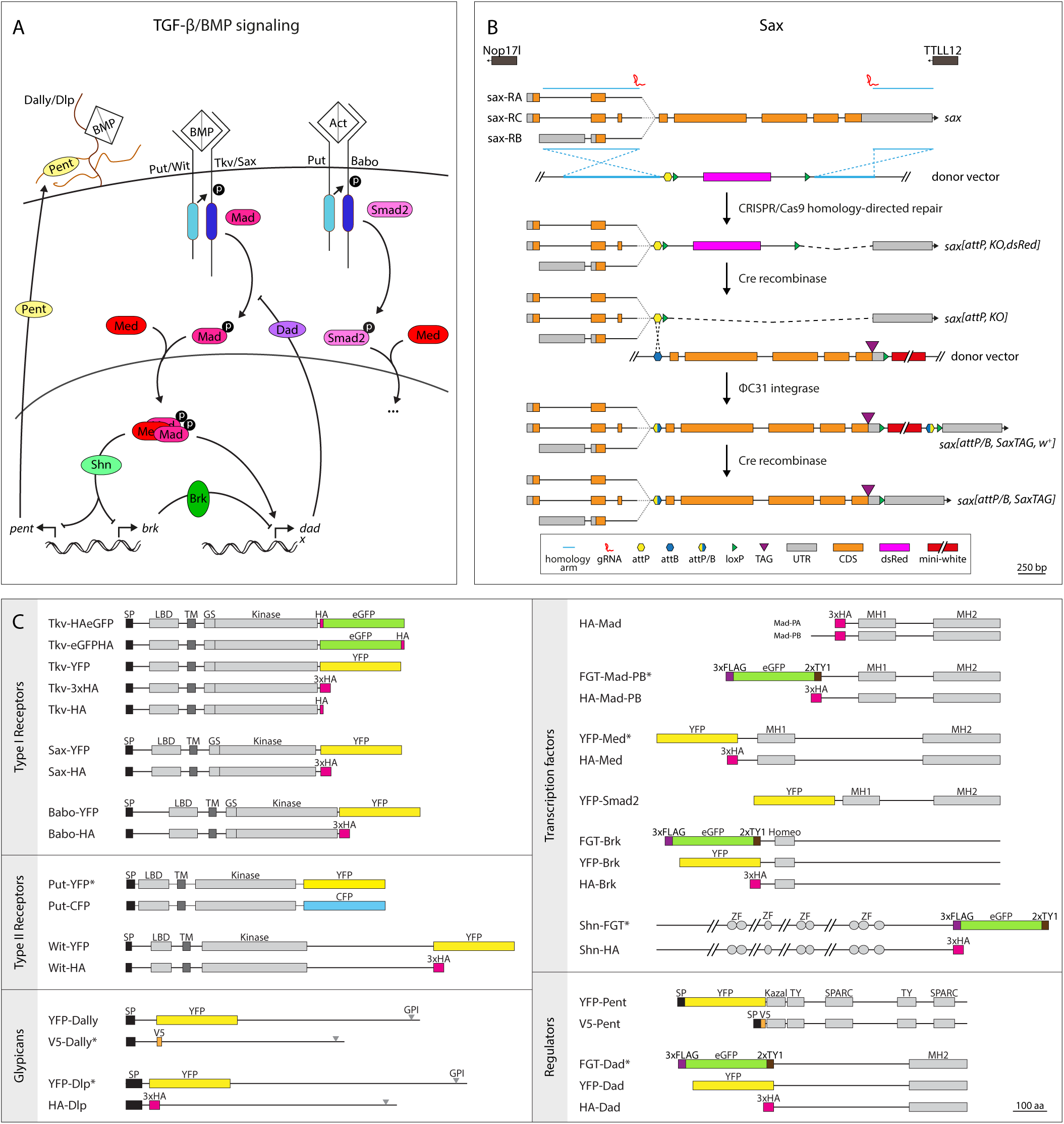
Genome engineering *Drosophila* TGF-β/BMP signaling. (A) Overview of the signaling pathway emphasizing on components that have been modified in this study (shown in color). (B) Two-step genome engineering strategy exemplified with Sax. CRISPR/Cas9 mediated homology-directed repair is used to replace the target sequence by an attP-containing cassette. After removal of the dsRed selection marker, the genomic locus is restored with tagged versions of the gene by standard ФC31 transgenesis. The mini-white selection marker is excised by Cre recombinase. Exact strategies for the other TGF-β/BMP components are depicted in Fig. S1. Note that in case of Dally, Pent, Tkv and Put homologous recombination as described in (Baena-Lopez et al., 2013) was used to generate the attP, KO lines. UTR = untranslated region, CDS = coding sequence, bp = base pairs. (C) In scale overview of tagged TGF-β/BMP components to visualize position of the tag (shown in color) in relation to protein domains. At least one but up to five tagged variants are available for all components. For simplicity, all components tagged with 3xHA are termed HA-X or X-HA. Note the exception of Tkv, for which the 3xHA tagged version is termed Tkv-3xHA, a single HA tagged version Tkv-HA and versions tagged with a single copy of HA and eGFP Tkv-HAeGFP and Tkv-eGFPHA. Variants marked by an asterisk are not viable in homozygosity. Grey rectangles at the glypicans indicate position of the GPI attachment site. SP = signal peptide, LBD = ligand binding domain, TM = transmembrane domain, GS = GS domain, Kinase = kinase domain, GPI = GPI anchor, MH1 = MH1 domain, MH2 = MH2 domain, Homeo = homeodomain, ZF = zinc finger domain, Kazal = Kazal domain, TY = thyroglobulin type 1 domain, SPARC = Secreted Protein Acidic and Rich in Cysteine domain, aa = amino acids.

### Assessing tissue and cellular distribution of tagged BMP components

With the above collection at hand, we tested whether our endogenously modified alleles can be used to monitor the distribution of the proteins in tissues and cells. We focused on three tissues, which critically depend on TGF-β/BMP signaling and its tight spatiotemporal regulation for proper development and function: the larval wing imaginal discs, the ovarian germarium and the developing egg chamber (Pyrowolakis et al., 2017; Upadhyay et al., 2017; Wilcockson et al., 2017). In the wing imaginal disc, both Activin and BMP signals provide growth (Activin) or growth and patterning (BMP ligands Dpp and Gbb) cues, with the latter acting as a morphogen. There is ample genetic and direct evidence that all modified genes of our library are expressed during larval wing development and are involved in the transmission and regulation of TGF-β/BMP signals. Indeed, we find robust expression of all tested components in the larval wing imaginal disc (Fig. 2, Fig. 3 and Fig. 4). Receptors (Tkv, Put, Sax, Babo and Wit) and glypicans (Dally and Dlp) can be readily visualized by immunostaining against the YFP and/or HA tags (Fig. 2B-H). In all cases, depletion of the corresponding transcripts either by gene-specific RNAi, or, in the case of YFP-tagged constructs, by the iGFPi system, resulted in a loss of signal confirming specificity (not shown). Our analyses indicate that most, if not all, receptors and glypicans display patterned distribution rather than being uniformly present in the wing disc epithelium. For the receptor Tkv, non-uniform expression along the anterior-posterior axis of the wing disc has been described, which depends on both BMP and Hedgehog (Hh) signaling that repress transcription of *tkv* in medial cells of the disc (Crickmore and Mann, 2006; Lecuit and Cohen, 1998; Tanimoto et al., 2000). The distribution of our Tkv variants is consistent with these observations (Fig. 2C). In addition, recent work has shown that the expression of Wit (a BMP type-II receptor) is transcriptionally activated by BMP signaling during larval wing development (Chayengia et al., 2019). These findings are recapitulated in the distribution of Wit-YFP, which is increased in medial and lower in lateral cells of the wing pouch (Fig. 2F). Put, the essential type II receptor for BMP signaling (Letsou et al., 1995; Ruberte et al., 1995), also displays modulated distribution along the anterior-posterior axis of the disc with lower levels at the positions of the two peaks of pMad on either side of the Dpp source (Fig. 2E). Sax, a type I receptor with a critical contribution in long range BMP signaling (Bangi and Wharton, 2006; Brummel et al., 1994; Haerry et al., 1998; Nellen et al., 1994; Nguyen et al., 1998; Penton et al., 1994; Xie et al., 1994), displays increased expression in lateral regions of the disc (Fig. 2B). At the same time Babo, a type I receptor for Activin controlling larval wing size (Brummel et al., 1999), is increased in medial cells (Fig. 2D).

**Figure 2.**
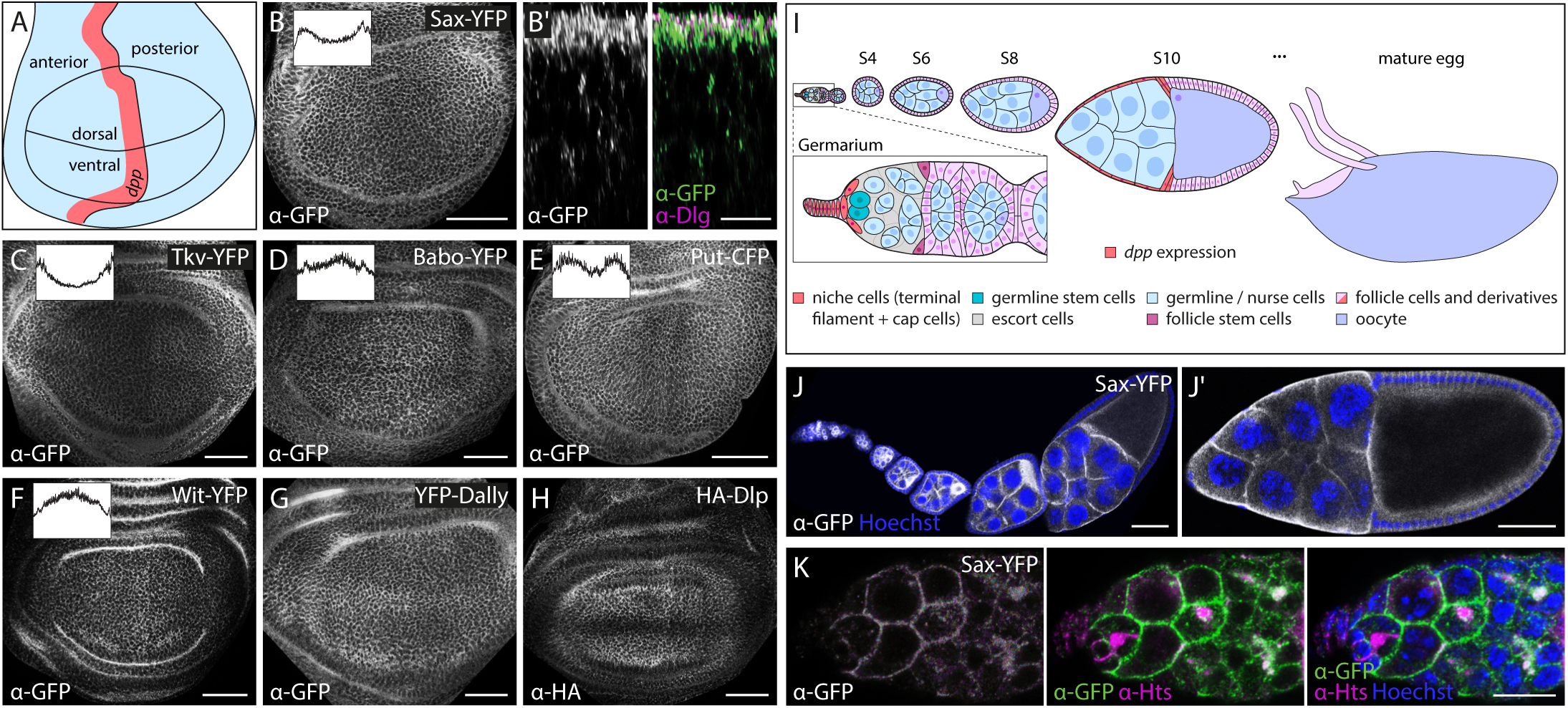
Tissue and cellular distribution of tagged TGF-β/BMP receptors and glypicans. (A) Schematic depiction of the larval wing imaginal disc showing the *dpp* expression domain in the anterior compartment in red. (B-H) Anti-GFP or anti-HA staining visualizes the distribution of receptors (B-F) and glypicans (G, H) in wing imaginal discs. Insets show intensity plots along the anterior-posterior disc axis. Scale bars: 50 μm. (B’) Subcellular localization of Sax-YFP in the disc proper in relation to the septate junction marker Discs large (Dlg) (magenta). Scale bar: 10 μm. (I) Schematic depiction of selected stages of oogenesis. Cells expressing *dpp* (niche cells in the germarium, stretched follicle cells in S10) are shown in red. S = stage. (J, K) Anti-GFP staining of Sax-YFP shows its distribution during oogenesis. Nuclei are visualized by Hoechst (blue) and spectrosomes by anti-hu li tai shao (Hts) staining (magenta). Scale bars: 50 μm (J), 10 μm (K).

**Figure 3.**
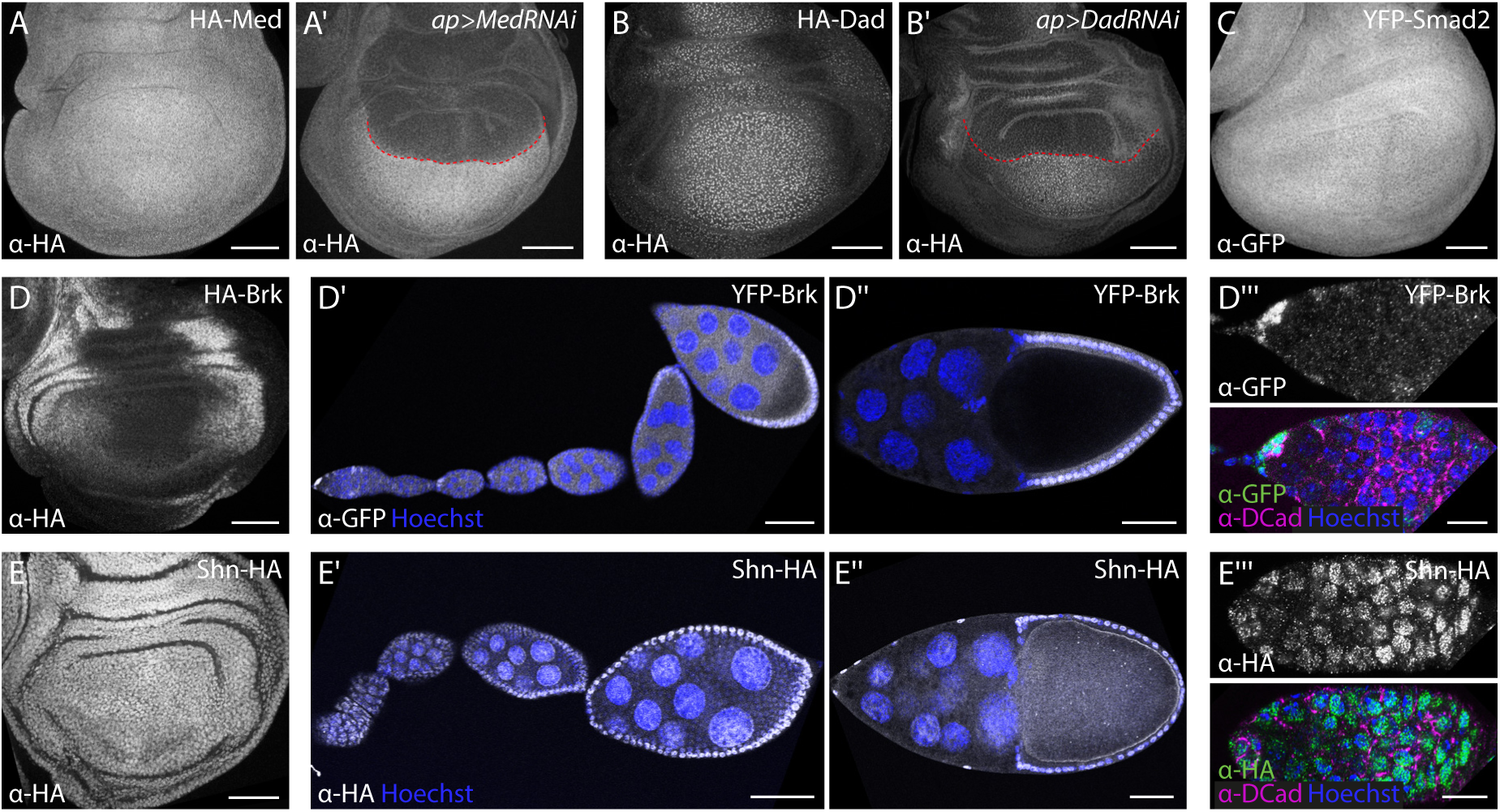
Tissue distribution of intracellular TGF-β/BMP components. (A-E) Anti-GFP or anti-HA staining visualizes the distribution of tagged Smad proteins (A-C) and transcription factors (D, E) in wing imaginal discs. (A’, B’) RNAi-mediated depletion of Med (A’) and Dad (B’) in the dorsal wing disc compartment using ap-Gal4 verifies specificity of the observed staining. Red dashed line indicates dorso-ventral compartment boundary. Scale bars: 50 μm. (D’, D’’’, E’, E’’’) Anti-GFP and HA-staining of YFP-Brk (D’, D’’’) and Shn-HA (E’, E’’’), respectively, show their distribution during oogenesis. Nuclei are visualized by Hoechst (blue) and DE-Cadherin (DCad) is shown in magenta. Scale bars: 50 μm (D’, D’’, E’, E’’), 10 μm (D’’’, E’’’).

**Figure 4.**
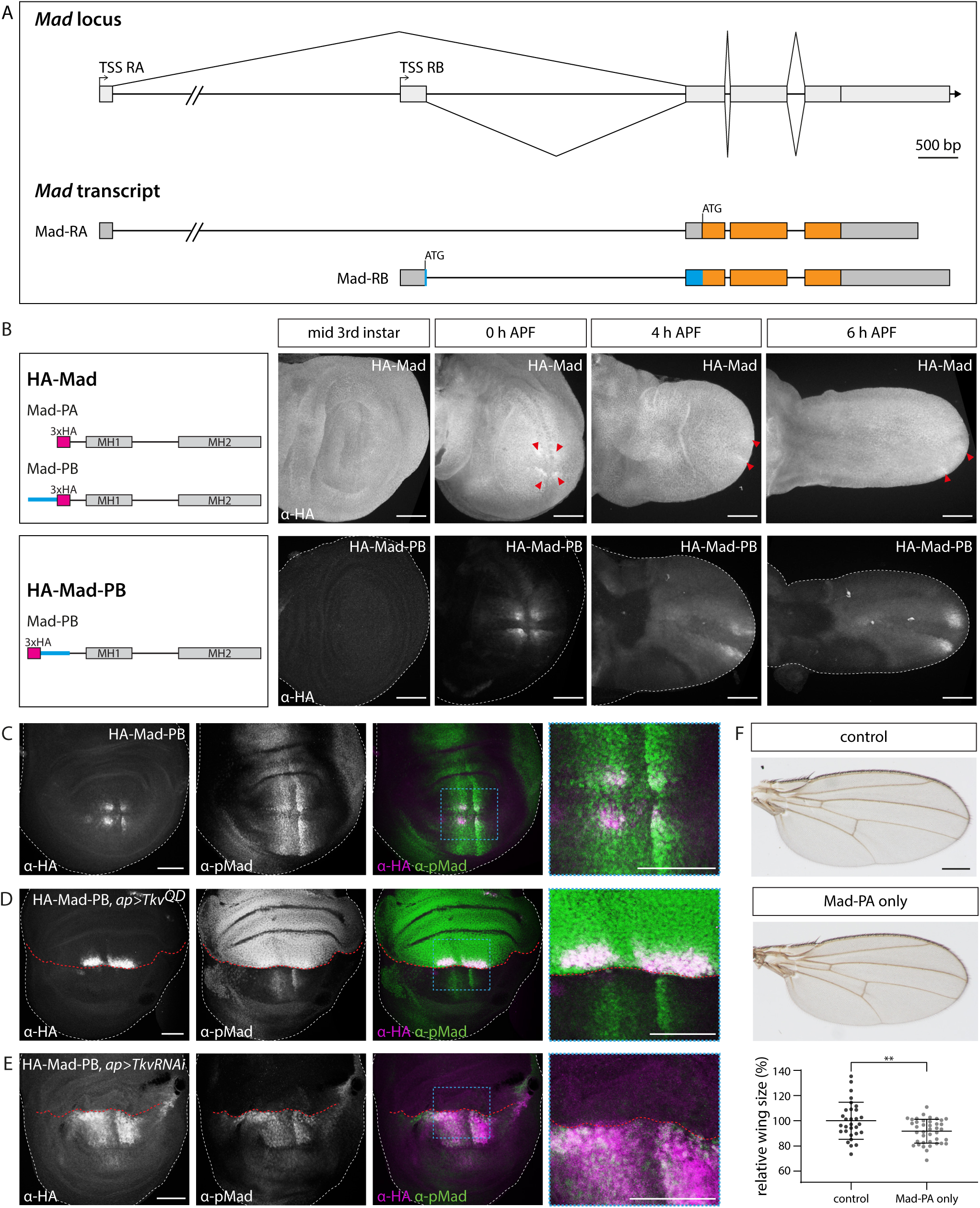
*Mad* isoform analysis. (A) *Mad* genomic locus and the predicted *Mad* transcript isoforms. Mad-RB contains a N-terminal extension (highlighted in blue). Orange color marks coding sequences and grey untranslated regions. TSS = transcriptional start site, bp = base pairs. (B) Anti-HA staining of endogenously tagged Mad versions visualized in mid 3^rd^ instar wing discs and early prepupal wings. HA-Mad tags both isoforms, while HA-Mad-PB exclusively tags Mad-PB. Discs are oriented anterior up and posterior down. Red arrowheads indicate patches of elevated protein levels, possibly corresponding to HA-Mad-PB in addition to the uniform HA-Mad-PA. APF = After puparium formation. Scale bars: 50 μm. (C-E) anti-HA and anti-pMad immunostaining in late larval wildtype wing discs (C) and discs expressing Tkv^QD^ (D) or Tkv-RNAi (E) in the dorsal compartment using ap-Gal4. Discs are oriented anterior left and posterior right. Red dashed line indicates dorso-ventral compartment boundary. Panels at the right are magnifications of the area indicated by the blue boxes. Scale bars: 50 μm. (F) Adult wings of control flies (carrying a reintegration of the wildtype genomic *Mad* sequence in *mad^[attP, KO]^*) and flies, which only contain Mad-PA. Graph shows relative wing size (%) of the indicated genotypes shown as mean ± SD. Dots represent measured size of individual wings (control: n = 30, Mad-PA only: n = 40). Statistical significance was analyzed by a two-tailed unpaired t-test with Welch’s correction assuming unequal variances. p≤0.01 = **. Scale bar: 250 μm.

The glypicans Dally and Dlp have been demonstrated to be modulated in their expression along the anterior-posterior or dorso-ventral axes of the wing and to differentially affect the activities of Wingless (Wg), Dpp, Hedgehog (Hh) and other signaling pathways during wing development (Baeg et al., 2001; Belenkaya et al., 2004; Franch-Marro et al., 2005; Fujise et al., 2003; Han et al., 2004; Han et al., 2005; Kirkpatrick et al., 2004; Kreuger et al., 2004; McGough et al., 2020; Simon et al., 2021; Simon et al., 2024). The complex expression pattern of Dally has been mainly deduced from enhancer trap lines and was shown to contribute to BMP gradient formation (Crickmore and Mann, 2007; de Navas et al., 2006; Fujise et al., 2001; Fujise et al., 2003; Makhijani et al., 2007). Broadly in line with these reports, our YFP-Dally displays increased protein levels in lateral regions and along the dorso-ventral compartment boundary of the wing disc while showing low expression in medial regions (Fig. 2G). However, our allele fails to reflect the prominent peak of *dally* expression in the *dpp* expressing stripe reported by *dally-lacZ* traps (Fujise et al., 2003). HA-Dlp, in agreement with previous reports (Han et al., 2005), is expressed at high levels in the dorsal and ventral cells of the pouch and is particular low in a broad stripe of cells straddling the dorso-ventral compartment boundary due to Wg-mediated repression (Fig. 2H).

We also addressed the distribution of the fusion proteins along the apical-basal axis of the polarized cells of the wing (Fig. 2B’ and Fig. S3). Apical and basolateral pools are detectable for all of the receptors and glypicans with the exception of Put, which seems to be excluded from apical cell membranes consistent with recent reports using UAS-based and rescue constructs of the gene (Peterson et al., 2022).

We observed robust but variegated expression of the tagged components during oogenesis (Fig. 2I-K and Fig. S4). Sax expression, for example, was mostly absent in follicle cells (FCs) but present in the germline with signals detectable already in germline stem cells (GSCs) and persisting in nurse cells and oocyte throughout oogenesis (Fig. 2J,K). This distribution is consistent with the requirements of Sax in the germline during oogenesis and egg chamber formation (Twombly et al., 1996; Xie and Spradling, 1998). Other receptors, including Tkv, Babo and Put, were expressed both in somatic and in germline cells, however at distinct spatial patterns and stages (Fig. S4A-F). Genetic studies have firmly established that Tkv transduces BMP signals in the GSC for stem cell maintenance and is also involved in the control of ligand distribution by patterned expression and expression in somatic cells of the ovarian niche (Luo et al., 2015; Ma and Xie, 2011; Michel et al., 2011; Tseng et al., 2018; Wilcockson and Ashe, 2019; Xia et al., 2012). Consistently, we observe widespread presence of Tkv in the germarium, with increased levels in the posterior-most cell population (Fig. S4B). In addition, Tkv is strongly expressed in the oocyte, where it displays a diffuse cytosolic distribution at early and a more juxtamembrane localization at later stages of oogenesis (Fig. S4A,A’). Moreover, and in agreement with its role in eggshell patterning, Tkv is found at membranes of follicle cells throughout oogenesis (Fig. S4A,A’) (Yakoby et al., 2008). Put, which is genetically required for GSC maintenance (Kawase et al., 2004), is present at membranes of germline cells throughout oogenesis and is also robustly expressed in follicle cells (Fig. S4E,E’,F). Babo, which has been genetically implicated in mediating Activin signaling in ovarian niche development (Lengil et al., 2015), is widely expressed in the germarium (but not in germline cells at later stages) and in follicle cells throughout oogenesis, a tissue for which no role of Activin/Babo signaling has been described so far (Fig. S4C,C’,D).

Both *Drosophila* glypicans have been functionally implicated in GSC niche homeostasis (Guo and Wang, 2009; Hayashi et al., 2009). Dally restricts BMP signaling to the anterior cells of the niche and is exclusively expressed in cap and terminal filament cells as judged by enhancer trap lines (Guo and Wang, 2009; Hayashi et al., 2009; Liu et al., 2010). Surprisingly, our tagged Dally allele, despite fully restoring fertility of *dally^[attP,KO]^* mutants, could hardly be detected in the germarium (Fig. S4G,H). The discrepancy to the reported *dally-lacZ* patterns might stem from low expression levels or limited stability of the protein in the cells of the niche. Indeed, at later stages of oogenesis, the same allele was readily detectable at membranes of developing follicle cells throughout oogenesis consistent with previous reports (Fig. S4G,G’) (Su et al., 2018). In contrast, Dlp was absent in the follicular epithelium at all stages of egg chamber development but clearly expressed in cells of the germarium. (Fig. S4I,I’,J). The distribution of Dlp in the germarium with the lower levels in escort cells (inner germariar cells) is in line with recent studies describing Dlp expression in the germarium and its role in regulating GSC differentiation (Tu et al., 2020; Waghmare et al., 2020).

Next, we assessed the distribution of intracellular components of the TGF-β/BMP signaling pathway in the same contexts (Fig. 3). The *Drosophila* Smads Med and Smad2 (for Mad see next section) were expressed at low and uniform levels in the wing imaginal disc, with no signs of nuclear enrichment (Fig. 3A,C). Dad, the single inhibitory Smad in *Drosophila*, was distributed in a pattern that is consistent with the well-established Dpp-dependent transcriptional regulation of the gene (Tsuneizumi et al., 1997; Weiss et al., 2010): *dad* transcription is directly activated by Smad complexes in medial cells of the disc and repressed by Brk in lateral cells (Fig. 3B). All signals were specific as they were lost upon RNAi-mediated depletion of the fusion proteins in the dorsal compartment (Fig. 3A’,B’). Tagged versions of Brk were distributed in a complementary nuclear gradient to the distribution of Dad, with highest levels in lateral cells and declining levels towards medial regions of the wing disc (Fig. 3D) (Marty et al., 2000; Müller et al., 2003; Pyrowolakis et al., 2004; Yao et al., 2008). Similarly, and in agreement with previous studies based on RNA *in situ* hybridization and reporter analyses (Charbonnier et al., 2015; Chen and Schüpbach, 2006; Shravage et al., 2007), the expression of tagged Brk in the follicular epithelium was restricted to posterior follicle cells that are devoid BMP signaling activity (Fig. 3D’,D’’). In both epithelia, *brk* expression is directly repressed by BMP signaling through Smad-dependent recruitment of Shn to *cis*-regulatory modules of the gene (Charbonnier et al., 2015; Marty et al., 2000; Müller et al., 2003; Pyrowolakis et al., 2004; Torres-Vazquez et al., 2000; Yao et al., 2008). Accordingly, Shn displays a uniform, nuclear expression pattern in both the wing disc and the follicular epithelium (Fig. 3E). In addition, Shn is present in the germarium and especially in the nuclei of germline cells, suggesting a role in BMP signaling mediated germline stem cell maintenance, which parallels findings from the male germline (Fig. 3’’’) (Matunis et al., 1997). Finally, Brk is weakly present in the germline but strongly expressed in somatic cells at the tip of the niche, consistent with a recent report on the role of Brk in regulating *dpp* expression in cap cells (Fig. 3D’’’) (Dunipace et al., 2022).

### An autoregulatory loop in *Mad* isoform expression

Mad is the founding member of the Smad transcription factor family and is the only BMP-responsive Smad encoded in the *Drosophila* genome. The ModEncode project predicts the existence of two *Mad* transcript isoforms (referred to as Mad-RA and Mad-RB respectively; see http://flybase.org/reports/FBgn0011648) arising by the use of alternative transcriptional start sites (Fig. 4A and Fig. S5). To our knowledge, Mad-RA (and the corresponding protein Mad-PA) is the isoform considered and exclusively used in previous studies that address Mad function mostly through UAS-based assays or cell culture expression. Mad-PB, the product of the Mad-RB transcript, bears a N-terminal extension of 70 amino acids that contains no discernable motifs and is of general low complexity (Fig. 4A,B and Fig. S5B). Based on developmental expression profiling data, Mad-RA transcripts are robustly detectable throughout development, while the Mad-RB specific exon is present during embryogenesis (but not maternally provided), declines during larval development and reappears at late larval stages and pupal development (Fig. S5A). We used our attP platform introduced into the *Mad* locus to address both the expression and function of the potential isoforms. We inserted three copies of the HA sequence directly after the ATG codons of either isoform to monitor protein distribution in wing imaginal discs. Note that, since Mad-PB represents a N-terminal, in-frame extension of Mad-PA, genomic insertion of the tag at the start codon of Mad-RA would theoretically result in a N-terminally HA-tagged Mad-PA and a Mad-PB version that carries an internal tag at amino acid position 71 (Fig. 4B and Fig. S5D). We refer to this construct as HA-Mad to indicate that both potential isoforms carry the HA tag. In contrast, insertion of the HA tag at the start codon of Mad-RB should exclusively label the Mad-PB polypeptide at its N-terminus; we refer to this allele as HA-Mad-PB. Larval wings at mid 3^rd^ instar stages display a ubiquitous HA-Mad distribution and some additional distinct patches of slightly increased levels in regions flanking the Dpp source cells are visible in late larval and prepupal stages (Fig. 4B, Fig. S5B). At the same time, HA-Mad-PB was absent at mid 3^rd^ instar wing discs but later appeared in four patches arranged around the intersection point of the compartment boundaries, which persisted and expanded during early pupal development (Fig. 4B, Fig. S5C). Notably, the patches of HA-Mad-PB coincided with peak levels of pMad in both the anterior and the posterior compartment, suggesting that BMP signaling is involved in the regulation of Mad expression (Fig. 4C). Indeed, increasing BMP activity by overexpression of a constitutive active Tkv receptor in the dorsal compartment resulted in an expansion of the two dorsal patches of HA-Mad-PB (Fig. 4D). Note that this manipulation also results in a drastic reduction of pMad and HA-Mad-PB in ventral cells. In reverse, RNAi-induced depletion of Tkv resulted in a loss of HA-Mad-PB expression in dorsal cells (Fig. 4E). At the same time, both the pMad gradient and HA-Mad-PB levels increased in ventral cells, with the effect being stronger in cells abutting the dorso-ventral compartment boundary. The non-autonomous effects in ventral cells most probably reflect redistribution of BMP ligands due to the increase or decrease of receptor levels in the dorsal compartment as reported before (for example see: Crickmore and Mann, 2006; Simon et al., 2024). Cumulatively, the results demonstrate that BMP signaling positively regulates Mad-PB expression.

To understand the relative contribution of the two isoforms in fly BMP signaling, we introduced mutations in the genomic sequence of *Mad* to express selectively Mad-PA or Mad-PB (Fig. S5E). We refrained from inserting epitope tags into these constructs to avoid confounding effects from such elements in the protein’s sequence. As a control, we also reintegrated the wildtype genomic *Mad* sequence using the same strategy in the *mad^[attP,KO]^* site. Flies carrying the control genomic sequence in homozygosity are viable and display no visible abnormalities, verifying the selected genome engineering approach (for wings see Fig. 4F “control”). As expected from the broad pattern of expression, flies devoid of the Mad-PA isoform are not viable. In contrast, flies able to express Mad-PA but not Mad-PB are viable and fertile. However, close examination of adult wings of such flies revealed a substantially smaller size than control wings, suggesting that Mad-PB contributes to final organ size (Fig. 4F). Taken together, our analyses suggest a regulatory loop in which BMP signaling, starting at late larval wing development, activates the expression of a distinct Mad isoform, which contributes to proper organ development.

### Membrane tethering, but not GPI-tethering, is essential for Dally**’**s function

We have recently used our engineering platforms to demonstrate that Dally, but not Dlp, is involved in the formation of the BMP signaling gradient (Simon et al., 2024). In addition, and in agreement with a recent study, we established that HS modification is required for Dally’s function in gradient formation (Nakato et al., 2024). Here we extend on these findings to interrogate another key structural feature of the protein: its membrane anchoring (Lin, 2004). Dally is tethered to the plasma membrane through a GPI anchor, however the exact function of membrane anchoring or the specific requirement of the lipid anchoring have not been addressed under conditions of physiological expression. We used our *dally^[attP,KO]^* platform to address these questions. Consistent with previous reports (Franch-Marro et al., 2005; Lin and Perrimon, 1999; Nakato et al., 1995), *dally* mutant (*dally^[attP,KO]^*) flies display a number of defects, including very low hatching rates, absence of male external genitalia and characteristic defects in wing development. Adult wings are smaller, with a “spitz” appearance (due to reduction of size along the anterior-posterior axis) and display distal truncations of longitudinal vein 5 (L5) (Fig. 5B,D,E and Fig. S1C). Reintegration of YFP-tagged Dally (YFP-Dally) resulted in flies that are fully viable and fertile (Simon et al., 2024). Wing discs of such flies display normal pMad distribution and develop into adult wings of normal size, proportions and pattern (Fig. 5 and Fig.S2). Compared to YFP-Dally, reintegration of Dally with a C-terminal truncation that removes the GPI-modification sites (YFP-Dally^ΔGPI^) resulted in a diffuse distribution in the disc epithelium consistent with the protein being secreted and not confined to plasma membranes (Fig. 5A). Similar to *dally^[attP,KO]^* flies, YFP-Dally^ΔGPI^ flies develop slowly and are semi-lethal with only few larvae developing into adult, sterile flies. In addition, the secreted form of Dally failed to restore pMad distribution, L5 formation and wing size of *dally* mutants (Fig. 5). Reestablishment of membrane anchoring by fusing a CD2 transmembrane domain to the C-terminus of Dally (YFP-DallyCD2), in contrast, reversed *dally* phenotypes. YFP-DallyCD2 flies were fully viable and fertile, displayed normal pMad distribution and adult wing morphology (Fig. 5). Cumulatively, the findings suggest that anchoring, but not necessarily GPI-anchoring, of Dally at the plasma membrane is important for Dally function.

**Figure 5.**
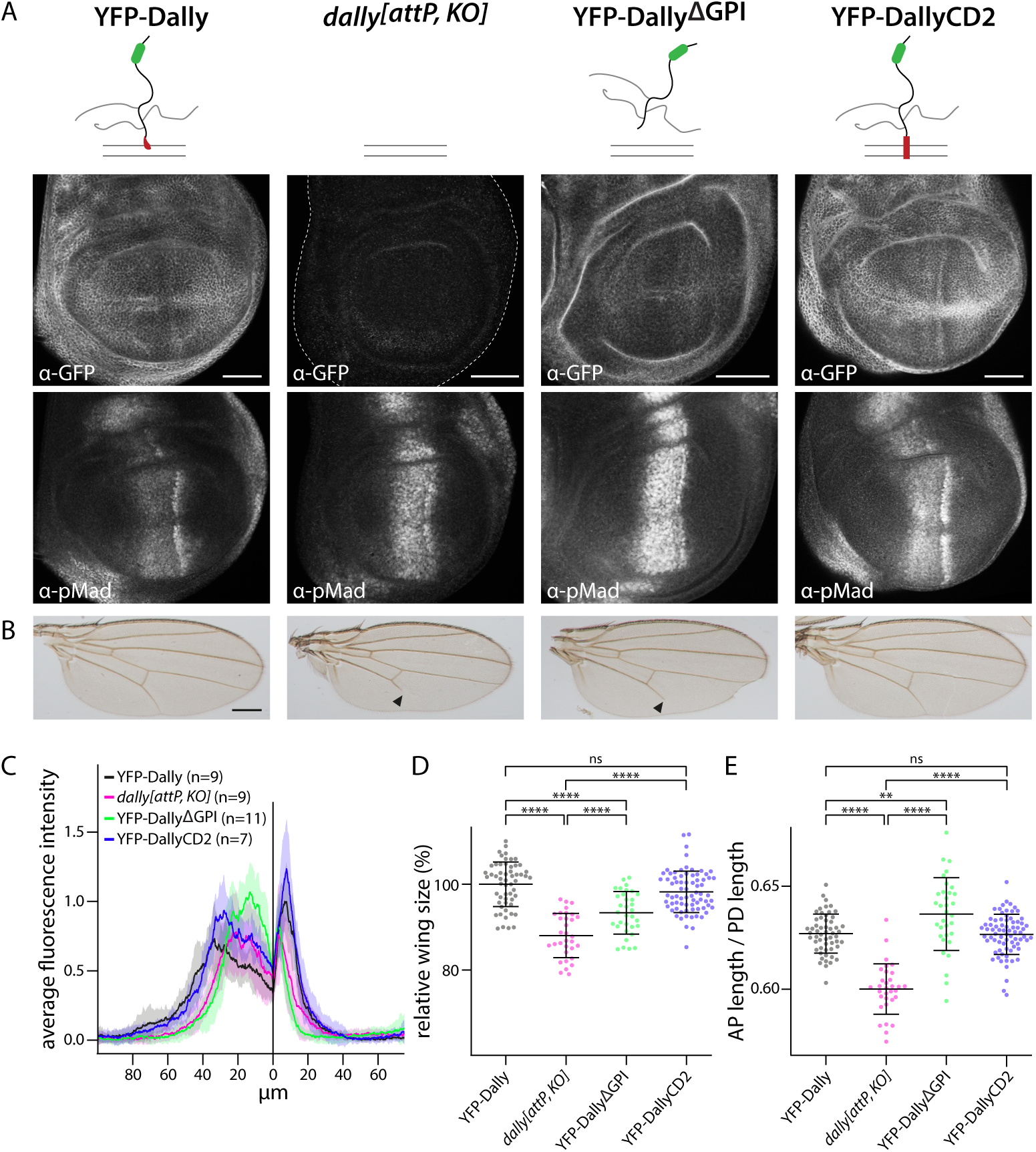
*In-locus* structure-function analysis of Dally. (A) Schematic illustration of analyzed versions of Dally: YFP-Dally, *dally^[attp,KO]^*, YFP-Dally^ΔGPI^ and YFP-DallyCD2. Anti-GFP and anti-pMad staining of wing discs of larvae carrying the modified Dally alleles in homozygosity. Scale bars: 50 μm. (B) Adult wings of the Dally versions shown in (A). All flies carry the respective modified Dally allele over the *dally^MH32^*null allele. Black arrowheads indicate truncation of longitudinal vein 5. Scale bar: 250 μm. (C) Average pMad fluorescence intensity of wing discs homozygous for the Dally versions indicated in A shown as mean ± SD. (D, E) Relative wing size (%) (D) and anterior-posterior (AP) length / proximal-distal (PD) length (E) of the indicated genotypes shown as mean ± SD. Dots represent measured values of individual wings (YFP-Dally: n = 56, *dally^[attp,KO]^*: n = 32, YFP-Dally^ΔGPI^ = 33, YFP-DallyCD2 = 76). All flies carry the respective modified Dally allele over the *dally^MH32^* null allele. Statistical significance was analyzed by a two-tailed unpaired t-test with Welch’s correction assuming unequal variances. p>0.05 = ns, p≤0.01 = ** and p≤0.0001 = ****.

### A complementary toolset for the manipulation of HA-tagged proteins

Recent studies have established a number of protein binder-based reagents that can be used to visualize and manipulate the activity of proteins that carry small peptide tags, including the HA epitope. Capitalizing on our collection of HA-tagged BMP components and inspired by a tool collection based on nanobodies against GFP and related fluorescent proteins (Caussinus et al., 2012; Harmansa et al., 2017; Matsuda et al., 2022), we generated and tested transgenic tools that are based on a single chain antibody specifically recognizing the HA peptide called Frankenbody (anti-HA-scFvX15F11 or FB) (Zhao et al., 2019). In addition to the FB-GFP and deGradHA, which we have shown recently to be useful for the visualization and degradation of proteins carrying single copies of the HA tag (Vigano et al., 2021), we have generated constructs that localize the FB to the outer (GrabHA_Ext_) or inner (GrabHA_Int_) surface of the plasma membrane, or at the basal lamina (GrabHA-ECM) (Fig. 6A). Such functionalized FB-fusions can be useful for trapping or enriching intracellular or extracellular proteins to the corresponding compartments. Indeed, GrabHA_Ext,_ expressed at the Dpp source efficiently trapped endogenously tagged HA-Dpp as shown by the compaction of the pMad gradient and the concomitant expansion of Brk expression (Fig. 6B). The effects were most prominent in the posterior compartment, where pMad was essentially absent and Brk was ectopically expressed in the whole compartment. The results, including the compartment asymmetry, are in line with a previous study using a different HA-trap version (Matsuda et al., 2021). We also found that our GrabHA_Int_ efficiently localized nuclear HA-tagged proteins to the plasma membrane while simultaneously depleting them from the nucleus. Expression of GrabHA_Int_ in the dorsal compartment of the wing disc resulted in nuclear depletion and strong accumulation of HA-Brk to the plasma membrane (Fig. 6C). Consistent with the role of Brk in reducing proliferation in the wing disc (Restrepo et al., 2014; Schwank et al., 2008), depletion of Brk from the nucleus resulted in a drastic increase of lateral proliferation rates in the dorsal compartment. Furthermore, expression of GrabHA_Int_ throughout the wing pouch induced a strong and characteristic adult wing overgrowth (Barrio and Milán, 2017). The effects on proliferation rates and adult wing morphology were comparable to the effects of iGFPi-mediated depletion of YFP-Brk (Neumüller et al., 2012) (Fig. 6C). In addition to the manipulation of HA-tagged proteins with the HA-based toolset, YFP-or FGT-fusion proteins of our library were also susceptible to the previously established GFP nanobody-based tools (Caussinus et al., 2012; Harmansa et al., 2017): Dorsal expression of deGradFP and morphotrap_Int_ resulted in efficient degradation and membrane translocation, respectively, of both nuclear Shn-FGT and cytosolic YFP-Smad2 (Fig. S6). Taken together, these results demonstrate that our library of modified TGF-β/BMP components is compatible with previously established and newly generated functionalized protein binder tools.

**Figure 6.**
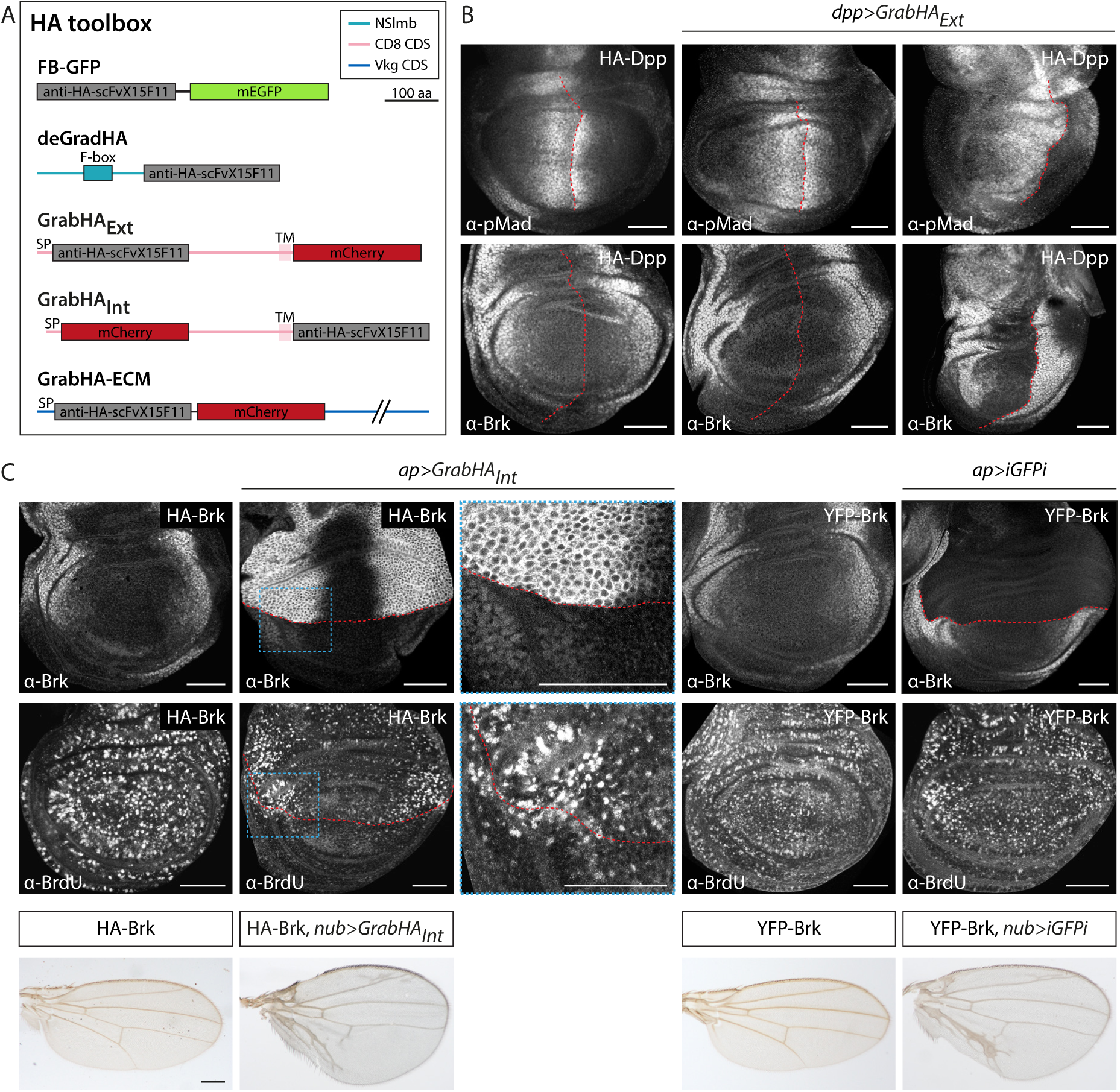
The HA toolbox. (A) Overview of the HA tools comprising FB-GFP, deGradHA, GrabHA_Ext_, GrabHA_Int_ and GrabHA-ECM. NSlmb = N-terminal part of Slmb, CDS = coding sequence, SP = signal peptide, TM = transmembrane domain, aa = amino acids. (B) anti-pMad and anti-Brk staining of wing discs expressing HA-Dpp, GrabHA_Ext_ in *dpp* producing cells or a combination of both. Anterior-posterior compartment boundary is marked by red dashed line. Scale bars: 50 μm. (C) Anti-Brk and anti-BrdU staining of wing discs expressing HA-or YFP-tagged versions of Brk either alone or in combination with the indicated tool in the dorsal compartment using ap-Gal4. Red dashed lines mark dorso-ventral compartment boundary. Adult wings carrying HA-or YFP-tagged versions of Brk either alone or in combination with overexpression of *GrabHA_Int_* or *iGFPi* throughout the wing pouch using nub-Gal4. Scale bars: 50 μm (wing discs), 250 μm (adult wings).

## Discussion

Here we present a comprehensive resource of genome engineered components of the core *Drosophila* TGF-β/BMP signaling pathway. Our experimental strategy provides two distinct libraries that we expect to be useful in the field. The first collection comprises attP-insertions in 14 genes of the signaling pathway, which on one hand can be used as molecularly defined null alleles, but also serve as a ready-to-use genome engineered platform for simple and efficient generation of variants of interest via ФC31-mediated integration. Our second collection capitalizes on this feature and consists of a number of functional, epitope-or fluorescently-tagged components, which are expressed from the corresponding endogenous genomic loci. We demonstrate that the generated tagged proteins can be used for capturing tissue and subcellular distribution in selected tissues. Our analysis of the tagged proteins does not only confirm expectations on the expression of the cognate genes but also highlights some new aspects. For example, the levels of the receptors of the pathway are modulated along the anterior-posterior axis of the larval wing, a feature that has been previously established only for Tkv. While we do not know whether this is a result of transcriptional or post-transcriptional regulation, these findings suggest that patterned expression of multiple receptors might contribute in shaping the BMP activity gradient in this tissue. Our analyses also highlight variegated and dynamic expression of most components of the pathway in eggshell patterning and in the germline stem cell niche.

Compared to recent efforts and advances in high-throughput epitope tagging, our gene-tailored approach, while not easily scalable, offers some advantages. Fosmid-based resources express epitope-tagged proteins from an additional gene copy that includes genomic regulatory regions (Sarov et al., 2016). While extremely useful, these lines tend to be restricted to genes of small genomic size and might result in abnormal expression patterns due to omission of regulatory elements. Indicatively, tagged versions of genes with large and complex genomic loci, such as *tkv*, *dally*, *dlp* or *pent* are not represented in current libraries. In addition, the current fosmid project only used C-terminal tagging, which might affect protein function in some cases. Furthermore, the additional presence of the endogenous, untagged gene is incompatible with certain experiments and would require a simultaneous genetic removal of the untagged allele. At the same time, gene tagging using protein trapping technologies is based on the insertion of artificial exon cassettes, thus excluding single-exon genes and drastically restricting the position of the tag within the protein sequence (Li-Kroeger et al., 2018; Nagarkar-Jaiswal et al., 2015; Venken et al., 2011). Our approach circumvents such problems and allows flexibility to specify the position of the tag for each gene individually. More important, the replacement of all or extensive sequence stretches of the coding regions by the attP-cassette allows for efficient reintegration of gene-variants, for example for structure-function analyses. In the present study, we used this feature to address isoform and structural requirements of Mad and Dally, respectively.

Our studies addressing isoform utilization of *Mad* established patterned expression of a previously uncharacterized isoform, Mad-PB, during wing development. Interestingly, the expression of Mad-PB is under positive control of BMP signaling, which activates an alternative promoter of the gene at late stages of larval development. In contrast to the essential Mad-PA isoform, flies lacking Mad-PB have no gross abnormalities or developmental delays. However, such flies have significantly smaller wings, suggesting that BMP-dependent activation of Mad-PB might be yet another regulatory loop in the system. At present, we can only speculate on the exact role of Mad-PB and its potential interaction with pMad. It is possible that BMP-dependent activation of Mad-PB boosts Mad levels during a critical late larval/early pupal stage, however there is no evidence that *Mad* expression levels are limited during wing development. In addition, the design of our “Mad-PA only” construct abolishes Mad-PB production at the protein level but still allows transcriptional activation of the corresponding transcript; since this isoform comprises Mad-PA, a general elevation of Mad levels should still be possible in these flies. Alternatively, Mad-PB, with its N-terminal extension, might regulate wing growth through the regulation of pMad. It is conceivable that in this scenario, Mad-PB, either directly or after phosphorylation through the activated receptors, regulates the activity of pMad in gene regulation, diverts pMad activity towards different targets, or even blunts pMad activity towards signal termination.

We also used our genomic platforms to address structural features of Dally, a glypican that regulates the BMP signaling gradient in the wing disc by mechanisms that are not yet fully understood. Dally, as all glypicans, is anchored to the plasma membrane by a GPI anchor, however the functional relevance of GPI-anchoring has not been directly addressed. GPI anchors are important for protein compartmentalization at membrane domains and are involved in the regulation of endocytosis and protein trafficking(Mayor and Riezman, 2004; Sezgin et al., 2017). These processes have already been implicated in BMP signaling activation and ligand dispersion in the wing disc and elsewhere (Akiyama et al., 2008; Belenkaya et al., 2004; Cai et al., 2019; Morawa et al., 2015; Norman et al., 2016; Romanova-Michailidi et al., 2021; Simon et al., 2024). In addition, a recent synthetic biology approach suggested that Dally might be directly involved in the transport of BMPs through repeating rounds of GPI-mediated detachment of BMP-loaded Dally and reinsertion into the membrane of neighboring cells (Stapornwongkul and Vincent, 2021; Stapornwongkul et al., 2020). Our results clearly demonstrate that membrane anchoring of Dally is indeed important for its function, as expression of a protein with a C-terminal truncation removing the GPI-modification sequences cannot rescue *dally* mutant phenotypes. However, restoring membrane tethering of the same construct by adding the transmembrane domain of the unrelated protein CD2 fully restores viability, fertility and wing patterning and growth. Thus, based on these findings, membrane tethering but not GPI-anchoring of Dally seems to be important for its function. Interestingly, it has also been suggested for Dlp, the second *Drosophila* glypican, to not require GPI-anchoring for the regulation of Wg signaling based on overexpression studies (Yan et al., 2009).

Finally, and capitalizing on our collection of epitope and fluorescently-tagged proteins, we tested whether our YFP/GFP- and HA-tagged proteins can be manipulated by recently established and newly generated functionalized protein binder tools. We demonstrate that YFP-, similar to GFP-tagged proteins, can indeed be efficiently recognized by the deGradFP and morphotrap_Int_ tools, resulting in degradation or trapping of the target protein, respectively (Caussinus et al., 2012; Harmansa and Affolter, 2018; Harmansa et al., 2017). Furthermore, our HA-tagged BMP components can also be destabilized by our recently established deGradHA tool, an anti-HA single chain antibody (Frankenbody, (Zhao et al., 2019)) fused to the F-box domain of Slmb, which channels proteins to ubiquitin/proteasomal degradation (Vigano et al., 2021). In addition, we established GrabHA_Ext_ and GrabHA_Int_, constructs, in which the Frankenbody is tethered to the plasma membrane facing either to the outside (GrabHA_Ext_) or to the cytosol (GrabHA_Int_), and demonstrated that they can indeed efficiently trap and localize extracellular proteins (HA-Dpp) or even miss-route and stabilize transcription factors at the plasma membrane (HA-Brk). We furthermore assume such tools to be functional when targeted to additional cellular domains and compartments.

In summary, we expect our collection of attP alleles as well as the library of epitope and fluorescently-tagged TGF-β/BMP components, which allow expression at physiological levels and endogenous expression pattern, to greatly assist research in BMP and Activin signaling in different *Drosophila* tissues and during various processes. In addition, our newly established functionalized HA-protein binder tools complement recent, similar collections and will certainly be useful for studies beyond BMP signaling (Harmansa et al., 2017; Kim et al., 2022; Xu et al., 2022).

## Material and methods

### Drosophila lines

Df(2L)Exel6011 (7497), Df(2R)Exel6054 (7536), Df(2R)BSC270 (23166), Df(3R)ED5644 (9090), Df(3L)Exel6099(7578), Df(3L)ED4414 (8702), Df(3L)ED4543 (8073), Df(2L)Exel7015 (7785), Df(3R)ED6361 (24143), Df(2R)Exel6060 (7542), Df(2R)X1 (1702), Df(3R)exel6176 (7655), Df(3R)BSC792 (27364), Dp(1;3)DC186 (30314), Dp(1;3)DC172 (30302), UAS-MadRNAi (31315), UAS-MedRNAi (31928), UAS-TkvRNAi (40937), UAS-GFPRNAi (iGFPi) (41556), ap-Gal4 (41245), Cre recombinase (851, 1092) and nos-Cas9 (54591, 78781, 78782) flies were provided by the Bloomington *Drosophila* Stock Center and UAS-DadRNAi (42840) by the Vienna *Drosophila* Resource Center. Other lines used were: vasa-ФC31 (Bischof et al., 2007), *dally^MH3^*^2^ (Franch-Marro et al., 2005), *pent^2^* (Vuilleumier et al., 2010), UAS_NSlmb-vhh-GFP4 (deGradFP) (Caussinus et al., 2012), UASTLOT_mCherry::CD8::vhh-GFP4 (morphotrap_Int_) (Harmansa et al., 2015), ap-Gal4, dpp-Gal4, nub-Gal4 (W. Gehring) and HA-Dpp (Bosch et al., 2017).

### Genome engineering and reintegration of tagged and/or modified gene versions

To replace specific parts of the genes encoding for the TGF-β/BMP components with attP cassettes, genome engineering based on ends-out homologous recombination (HR) or CRISPR-induced homology directed repair (HDR) was used. To manipulate *dally*, *pent*, *tkv* or *put*, homology arms flanking the targeted region of the respective gene were inserted into the pTV^Cherry^ targeting vector (DGRC #1338) and the constructs were further integrated, mobilized as well as linearized as described (Baena-Lopez et al., 2013). Modified progeny with successful HR-mediated integration of the attP cassette was identified by the red eye color of the mini-white containing selection cassette, which was subsequently removed using Cre recombination. For all other components, CRISPR-based HDR was applied. Guide RNA target sites flanking the region of interest were selected using the publicly available tools FlyCRISPR Optimal Target Finder (Gratz et al., 2014) and DRSC Find CRISPRs, Harvard Medical School (https://www.flyrnai.org/crispr/) and the genomic sequence at the target site was validated by sequencing. PCR-amplified homology arms were cloned into the pHD-dsRed-attP reintegration vector (Addgene #51019 (Gratz et al., 2014)) and guide RNAs were cloned into plasmid pCFD3-dU6:3 (for Dlp, Mad and Med) or pCFD4-U6:1_U6:3 (for Babo, Brk, Dad, Sax, Shn, Smad2 and Wit) (Addgene #49410 and #49411, (Port et al., 2014)). Co-injection of the homology arms and guide RNA containing plasmids into *nos-Cas9* expressing flies resulted in HDR and successful integration of the attP cassette was identified by expression of the 3xP3-dsRed marker, which was then excised using Cre recombination. Primers used for amplification of the homology arms as well as the generation of the guide RNA plasmids are listed in Table S1 and deletion strategies for all components are schematically depicted in Fig. 1B and S1A. Introduced deletions were verified by sequencing (Fig. S1B).

In the second step, standard ФC31/attB integration was used to reintegrate either tagged and/or modified gene versions (e.g. for Mad and Dally) into the attP sites using the RIVwhite vector (DGRC #1330 (Baena-Lopez et al., 2013)). Depending on gene architecture and the introduced deletion, generated reintegration vectors contained either the deleted sequence or full-length cDNA. Primers used for the generation of the reintegration vectors are listed in Table S2 and schematic depictions of the reintegration strategies can be found in Fig. 1B and S1A. All generated constructs were verified by sequencing.

Some of the generated attP,KO as well as some reintegration lines were previously published: *tkv^[attP, KO]^*, Tkv-3xHA, *pent^[attP, KO]^*and YFP-Pent (Norman et al., 2016; Tracy Cai et al., 2019), Tkv-HAeGFP (Vigano et al., 2021), *brk^[attP, KO]^*, HA-Brk, *shn^[attP, KO]^* and Shn-HA (Vuilleumier et al., 2022), dally*^[attP, KO]^*, YFP-Dally, dlp*^[attP, KO]^*and HA-Dlp (Simon et al., 2024).

### Generation of HA toolbox

pUASTLOTattB_anti-HA_fb_GFP (referred to as FB-GFP) and pUASTLOTattB_deGradHA (referred to as deGradHA) have been previously described (Vigano et al., 2021). pUASTLOTattB_GrabHA_Ext_ and pUASTLOTattB-GrabHA_Int_ were generated by replacing vhh-GFP4 of pUASTLOTattB_vhh-GFP4::CD8::mCherry (Addgene #163917, (Harmansa et al., 2015) or pUASTLOTattB_mCherry::CD8::vhh-GFP4 (Addgene #163930, (Harmansa et al., 2017), respectively, with the PCR-amplified Frankenbody anti-HA-scFvX15F11_mEGFP (referred to as anti-HA_fb) (Vigano et al., 2021; Zhao et al., 2019). For pUASTLOTattB_GrabHA-ECM, vhh-GFP4 was cut out of pUASTLOTattB_vhh-GFP4::Vkg::mCherry (Addgene #163929, (Harmansa et al., 2017) and anti-HA_fb was integrated. All generated constructs were verified by sequencing. Flies carrying UASTLOT_anti-HA_fb_GFP, UASTLOT_deGradHA, UASTLOT_GrabHA_Ext_, UASTLOT_GrabHA_Int_ or UASTLOT_GrabHA-ECM on chromosome 3L at position 68A4 (attP2) were generated by standard ФC31/attB transgenesis. Primers used for the HA toolbox plasmids are listed in Table S3.

### Dissection and antibody staining of *Drosophila* tissues

For larval wing imaginal disc immunostaining, 3^rd^ instar larvae were collected in Phosphate Buffered Saline (PBS), cut into half, the posterior end was discarded and the remaining anterior part was inverted. The carcasses were roughly cleaned from excessive tissue and fixed in 4 % Paraformaldehyde (PFA) for 20 minutes. Carcasses were rinsed twice in PBSTx (0.1 % Triton X-100 in PBS), washed two times for 10 minutes in PBSTx and incubated for 1 hour in blocking solution (5 % NGS in PBSTx). Samples were then incubated with primary antibodies in blocking solution over night at 4 °C. On the next day, carcasses were rinsed twice, washed three times for 20 minutes in PBSTx and incubated with fluorescently labelled secondary antibodies and Hoechst 33342 in blocking solution for at least 2 hours. Samples were rinsed and washed three times for 10 minutes in PBSTx. After rinsing with PBS, the wing discs were fine-dissected and mounted in VECTASHIELD® Antifade Mounting Medium (Biozol).

For prepupal wings, white prepupae (0 hours APF) were collected and incubated at 25 °C until the desired age. Wings were then isolated from the prepupae, transferred to 4 % PFA, fixed for 20 minutes and immunostained as described above.

Ovaries were isolated from 2-to 4-day old female flies in PBS and were carefully opened to separate individual ovarioles, which were then fixed and stained as described for the wing discs.

BrdU labeling of larval wing imaginal discs was performed as follows: 3^rd^ instar larvae were dissected as described above, transferred into S2 medium and BrdU (0.1 mg/ml) was added to the medium. After 15 minutes incubation on a rocking platform, carcasses were fixed in 4 % PFA for 20 minutes, rinsed three times in PBSTx and incubated 45 minutes in 2N hydrochloric acid. After two short incubations with Na_3_BO_3_ (0.1 M, pH 8.5) for 2 minutes, samples were rinsed and washed twice in PBSTx for 10 minutes. A 20-minute incubation in blocking solution was followed by incubation with an anti-Brdu antibody over night at 4 °C. On the next day, samples were rinsed twice, washed three times for 10 minutes with PBSTx and blocked for 20 minutes in blocking solution. Fluorescently labelled secondary antibodies and Hoechst 33342 in blocking solution were added for at least 2 hours. Further steps after secondary antibody incubation were performed as described above.

The following antibodies were used in this study: chicken anti-GFP (1:1000, Abcam), rat anti-HA (1:200, Roche), mouse anti-Dlg (1:50, DSHB), rabbit anti-pMad (1:500, Abcam), rabbit anti-pMad (1:500, Cell Signalling), guinea pig anti-Brk (1:500, Hilary Ashe), mouse anti-Wg (1:40, DSHB), mouse anti-Ptc (1:40, DSHB), rat anti-DCad (1:50, DSHB), mouse anti-Hts (1:10, DSHB), mouse anti-Brdu antibody (1:100, BD Bioscience), Alexa fluorophore-conjugated secondary antibodies (1:500; Thermo Fisher Scientific) and Hoechst 33342 (1:5000; Invitrogen).

### Imaging and image processing

All immunostained samples were imaged with a ZEISS LSM 880 laser scanning confocal microscope (Life Imaging Center (LIC), Hilde Mangold Haus, University of Freiburg), which provides the additional opportunity to use a Airyscan detector or the FAST Airyscan mode for image acquisition. Whenever Airyscan detection was used, raw data was processed using the Airyscan processing function in the ZEISS ZEN Black 2.3 software. Images were analyzed and processed with Fiji and Adobe Photoshop. Figures were prepared using Adobe Illustrator.

### Quantification of pMad intensity

To obtain average intensity profiles of different samples of one genotype, first average intensity z projections of three consecutive z slices were generated using Fiji and signal intensity profiles along the AP wing disc axis were obtained in the dorsal compartment using identical sized boxes and the plot profile function in Fiji. The measured values were transferred to Excel (Microsoft) and values of different samples of the same genotype were aligned along the AP compartment boundary based on Ptc and/or pMad stainings and an average intensity profile was generated using the script *wing_disc-alignment.py* (Simon et al., 2024). Average intensity curves of different genotypes were then aligned and compared using the script *wingdisc_comparison.py* (Simon et al., 2024), normalizing the data with the smallest value of each experimental condition (normalization option *"n"*). The figure of the average intensity profile plots containing the SD for all compared genotypes was then prepared using Adobe Illustrator.

### Adult wing preparation, imaging and quantification

Adult flies were collected in isopropanol and dissected wings were mounted in Euparal mounting medium on a glass slide and covered with a cover slip. Images were acquired on a Leica MZ Apo using a ZEISS Axiocam 305 color camera.

Wing area and/or AP and PD length were measured using Fiji or Adobe Illustrator and values were transferred to Excel (Microsoft). Plots were generated using Prism (GraphPad) and adapted in Adobe Illustrator. Statistical significance was analyzed by a two-tailed unpaired t-test with Welch’s correction assuming unequal variances using Prism (GraphPad) with p>0.05 = ns, p≤0.05 = *, p≤0.01 = **, p≤0.001 = *** and p≤0.0001 = ****.

## Acknowledgements

We thank Jean-Paul Vincent, Markus Affolter, Hilary Ashe and Konrad Basler for fly lines, antibodies and plasmids. Stocks obtained from the Bloomington *Drosophila* Stock Center (NIH P40OD018537) were used in this study. We thank the Developmental Studies Hybridoma Bank (DSHB, The University of Iowa, Department of Biology) for monoclonal antibodies and the staff of the Life Imaging Center (LIC) of the University of Freiburg for help with microscopy resources and the excellent support in image recording and analysis. We are grateful to Elizabet Savkova for help with fly crosses and wing preparations, Thorina Boenke and Mark Norman for input at early stages of the project and Daniel Armbruster for critical comments on the manuscript and help with running the script for the quantification of pMad. Finally, we are indebted to Annette Neubüser for continuous support and to Daniela Reuter-Schmitt for excellent technical assistance.

## Funding

This work was funded by the Deutsche Forschungsgemeinschaft (DFG, German Research Foundation) under Germany’s Excellence Strategy EXC-2189-Project ID 390939984 and PY72/2-1.

---

**Figure S1.**
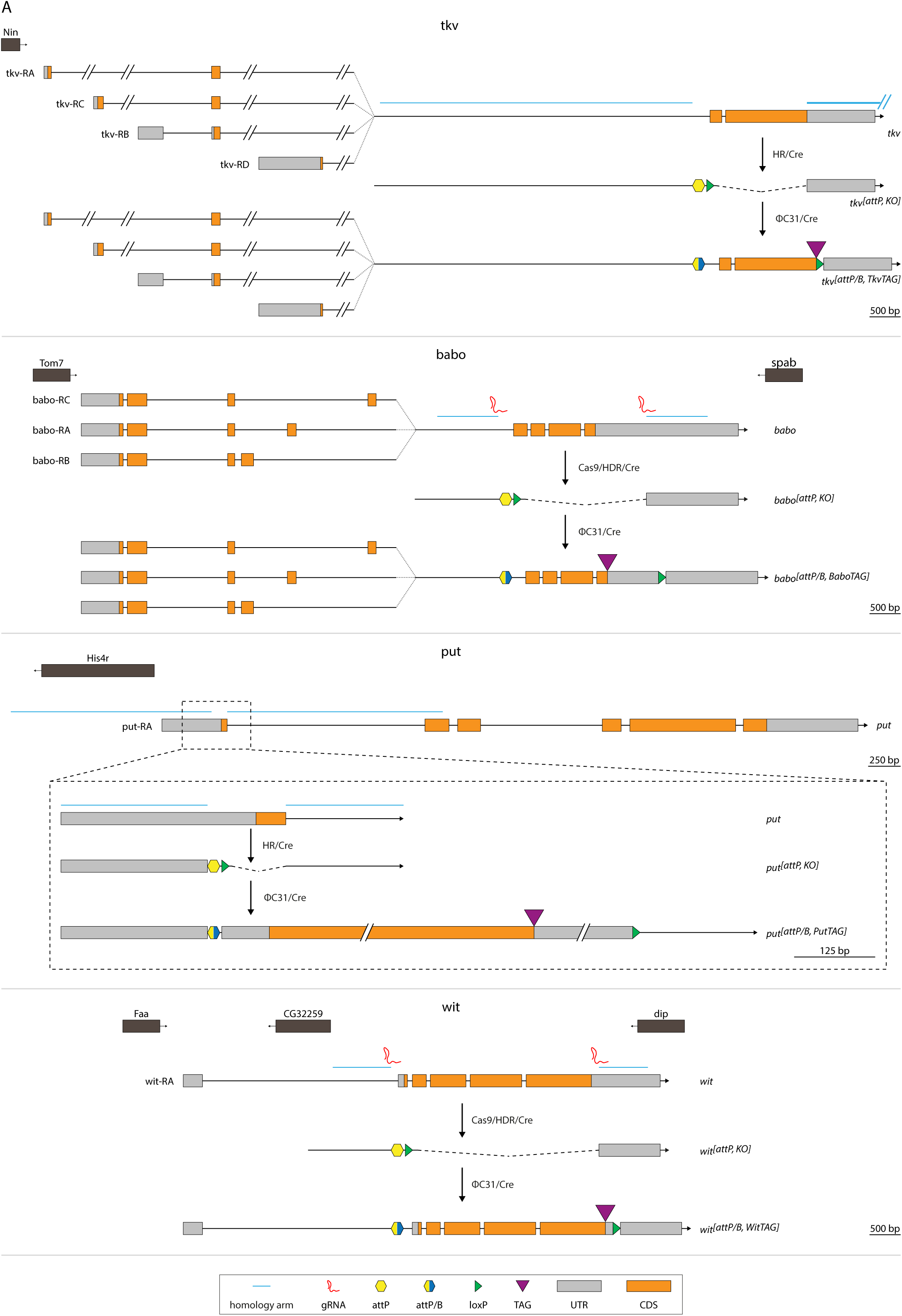
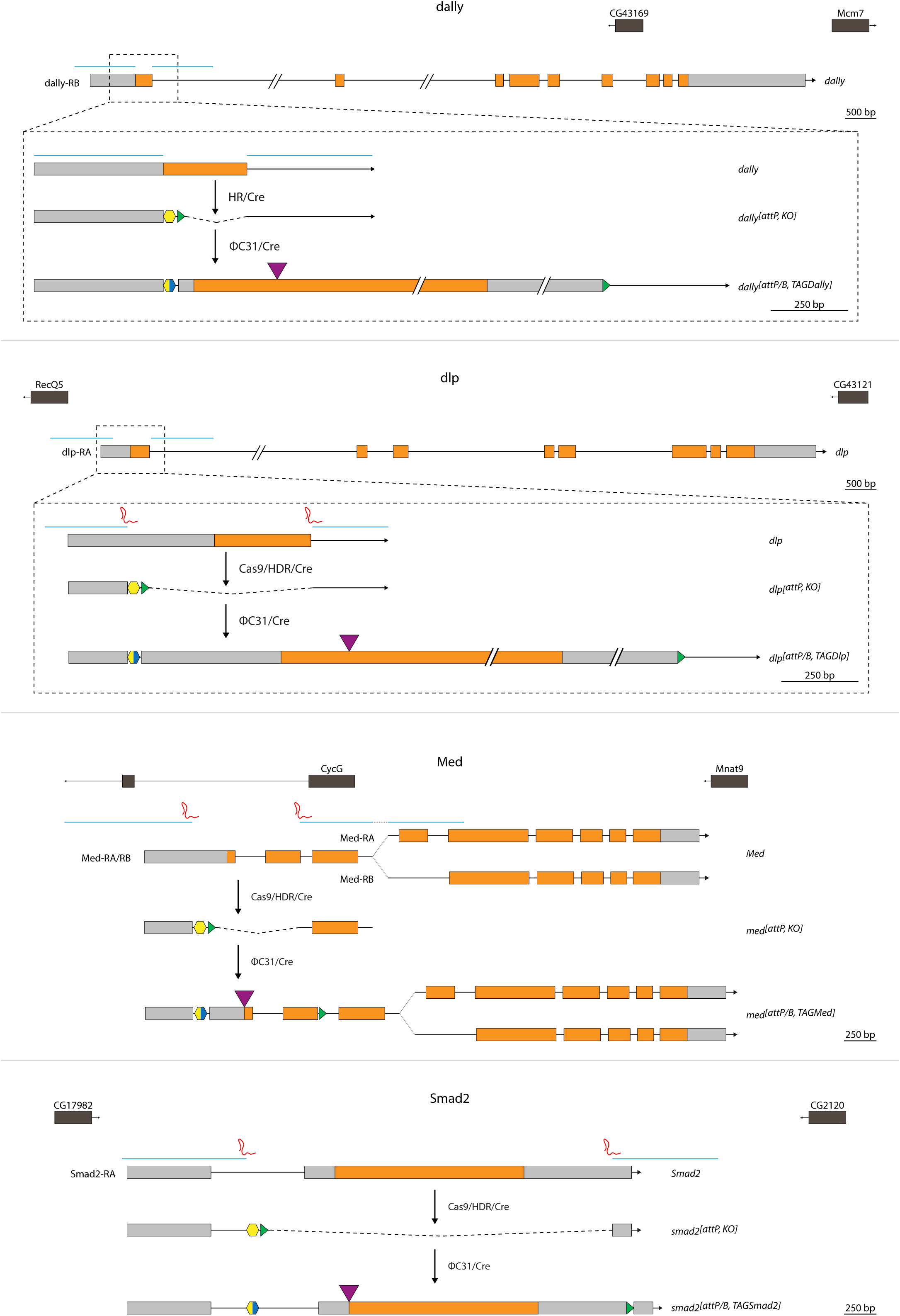
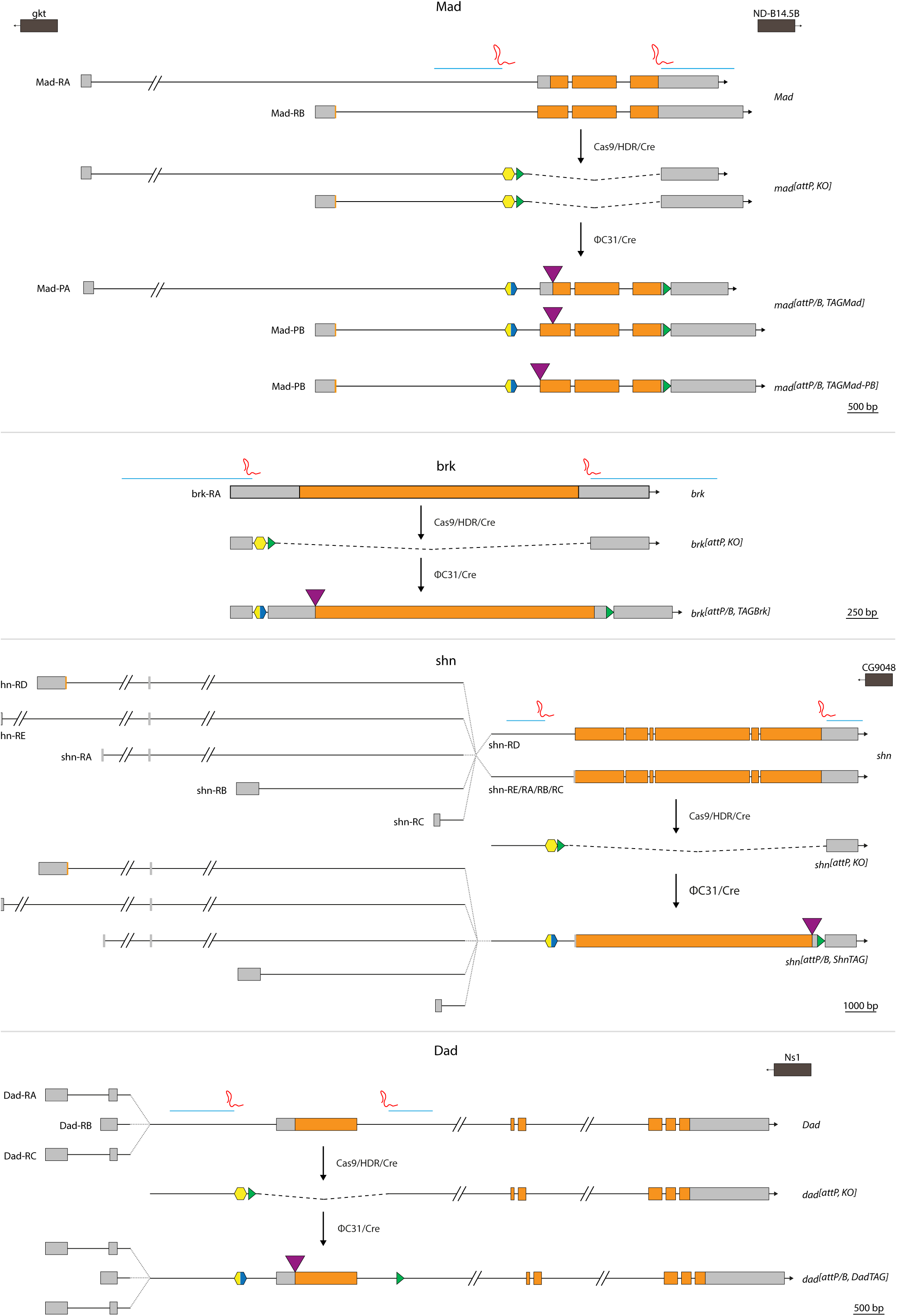
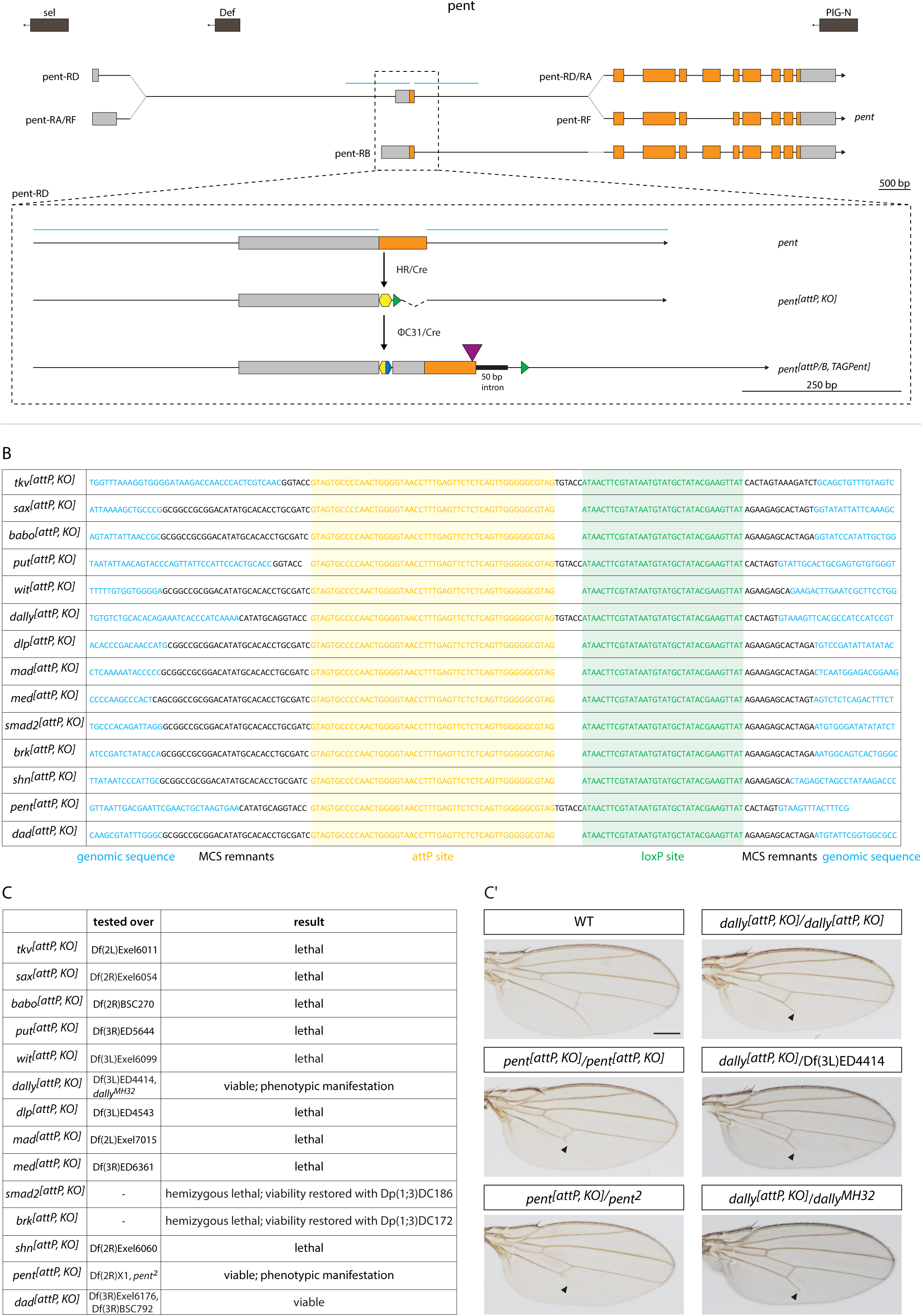
Genome engineering of components of the TGF-β/BMP signaling pathway. (A) Simplified overview of two-step genome engineering strategies for all modified TGF-β/BMP components. Intermediate steps bevor removal of selection markers are not shown (compare Fig. 1B). Note that in case of Dally, Pent, Tkv and Put homologous recombination was performed as described in (Baena-Lopez et al., 2013) instead of CRISPR/Cas9 based homology-directed repair to generate the attP, KO lines. Color code as shown in box under *wit* strategy. HR = homologous recombination, HDR = homology-directed repair, UTR = untranslated region, CDS = coding sequence, bp = base pairs. (B) Exact sequences of modified loci after integration of the attP site and removal of the selection marker. Color code: blue = genomic sequence, yellow = attP site, green = loxP site, black = remnants of multiple cloning site (MCS). (C) Generated knockout alleles were genetically verified by testing over the indicated fly lines (chromosomal deficiencies, previously described null alleles or mutants, chromosomal duplications for genes on X chromosome). *dally^[attP, KO]^* and *pent^[attP, KO]^* are viable over the tested chromosomes and show wing phenotypes as shown in C’. Wings are smaller than wild type (WT) wings and show truncations of longitudinal vein 5 (indicated by black arrowheads). Scale bar: 250 μm.

**Figure S2.**
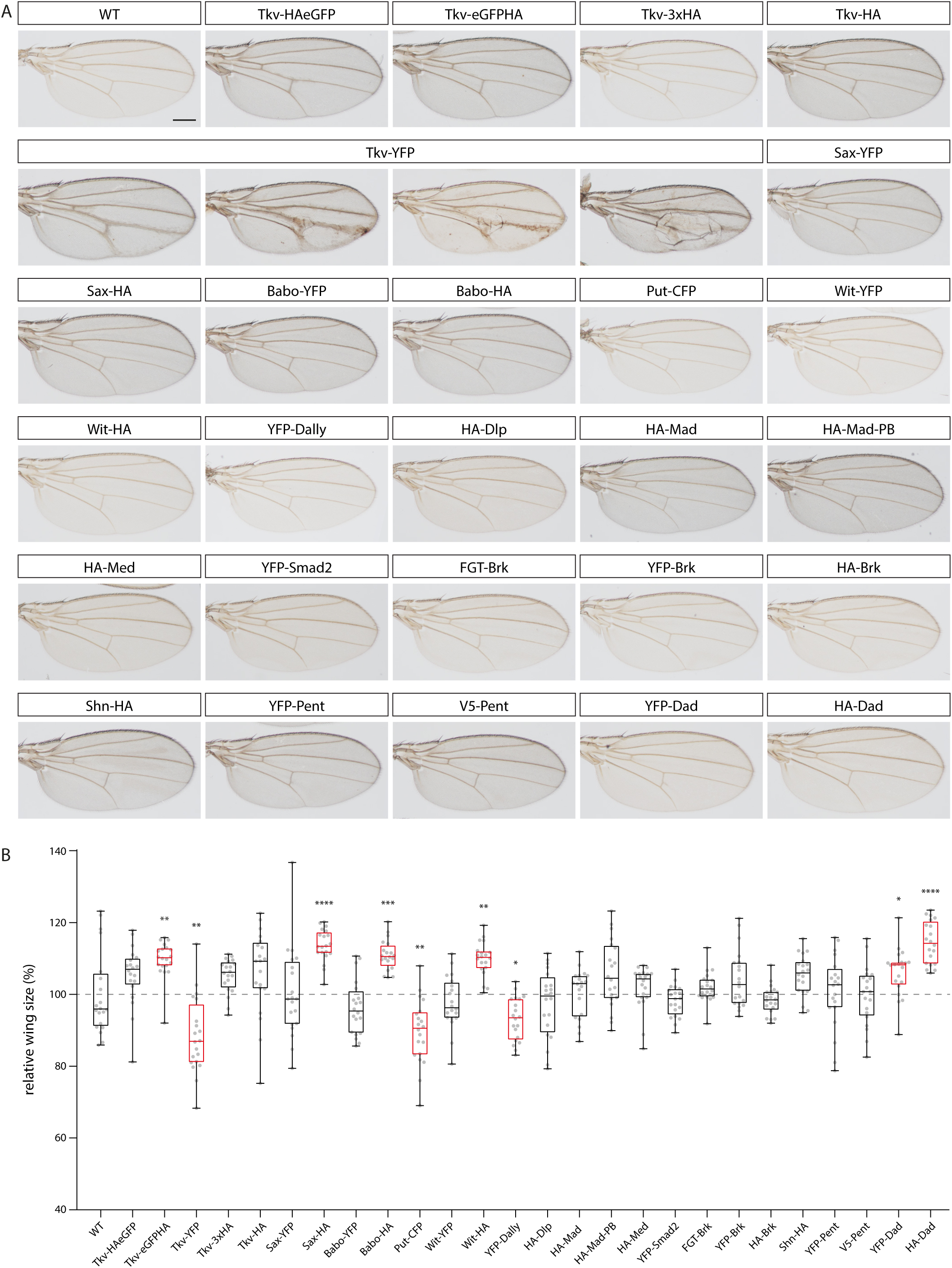
*In-locus* modified TGF-β/BMP components support wing development. (A) Adult wings of wildtype (WT) flies and flies carrying the tagged components in homozygosity. All wings show normal morphology and patterning except for wings of flies homozygous for Tkv-YFP, which frequently show thickening of veins and blisters. Scale bar: 250 μm. (B) Graph shows distribution of relative wing size (%) as boxplot for the indicated components. Dots represent measured size of individual wings (n = 20 for all). Statistical significance was analyzed by a two-tailed unpaired t-test with Welch’s correction assuming unequal variances comparing the wing size of the tagged components to WT. Instances deviating significantly from the control are highlighted in red. p>0.05 = ns (not labelled in the graph), p≤0.05 = *, p≤0.01 = **, p≤0.001 = *** and p≤0.0001 = ****.

**Figure S3.**
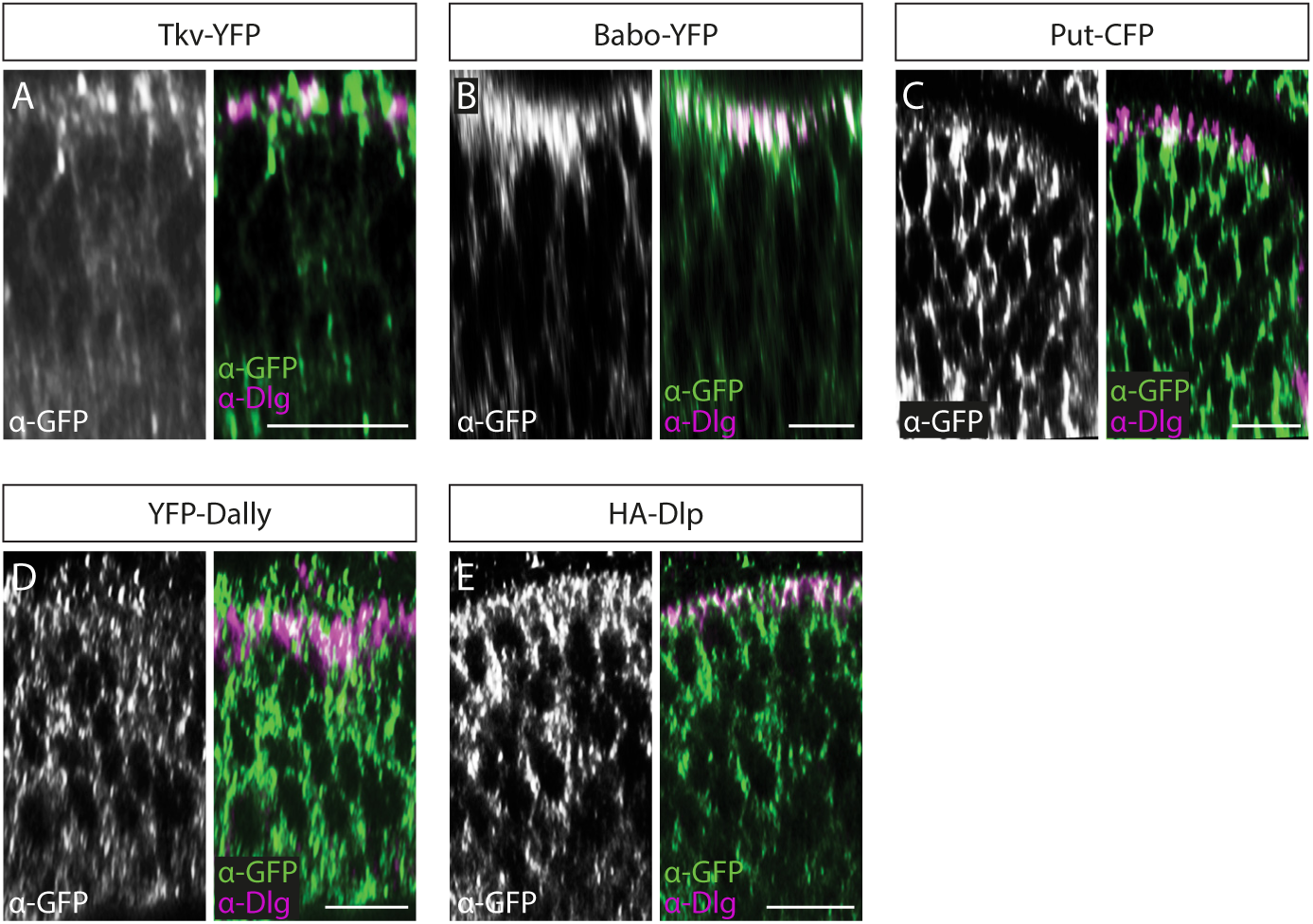
Subcellular localization of tagged receptors and glypicans. (A-E) Distribution of YFP-, CFP-or HA-tagged components in the disc proper visualized by anti-GFP or anti-HA staining in relation to Discs large (Dlg) (magenta). Scale bar: 10 μm.

**Figure S4.**
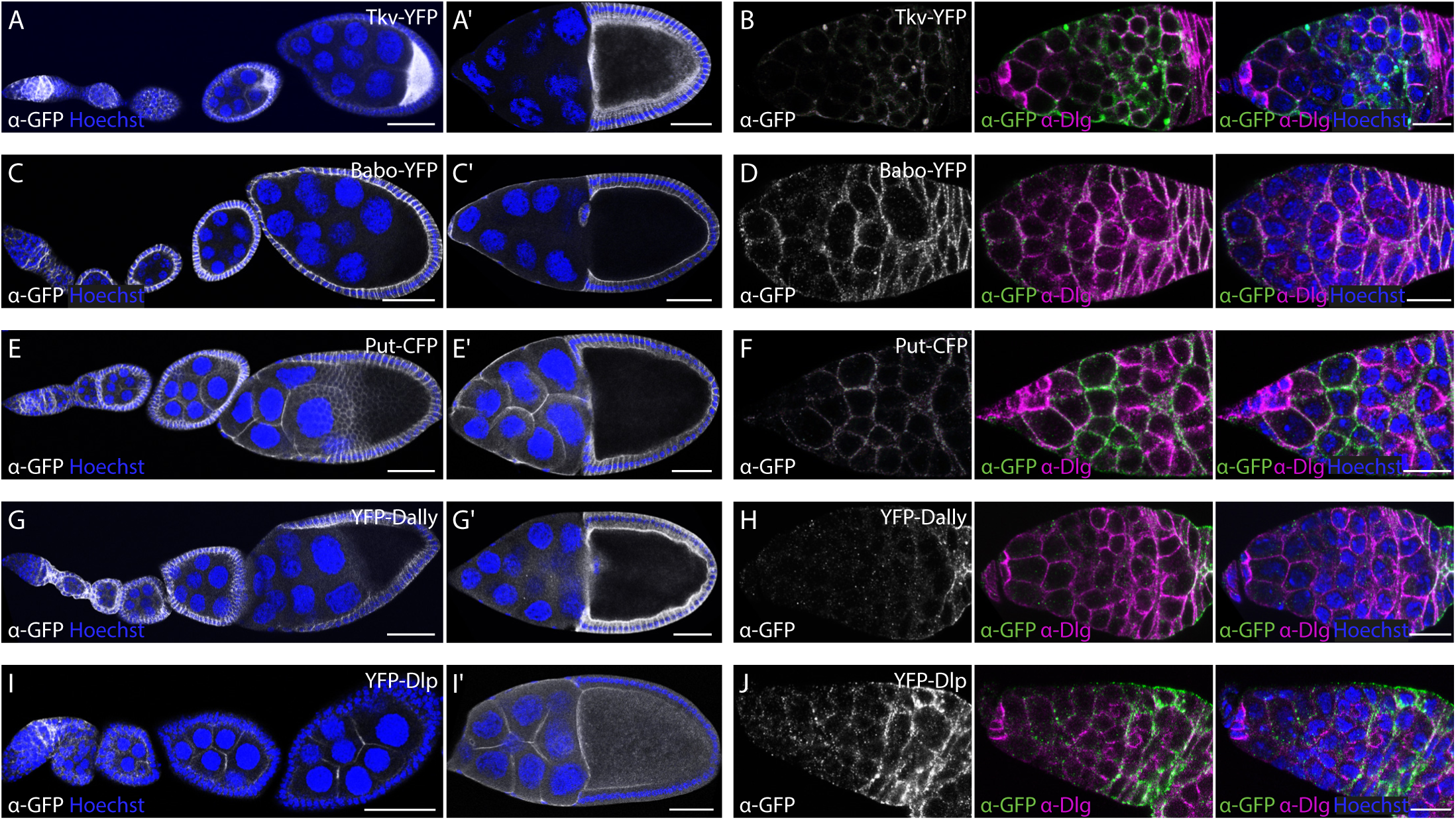
Distribution of tagged receptors and glypicans during oogenesis. (A-J) Anti-GFP staining of YFP-or CFP-tagged components shows their distribution during oogenesis. Nuclei are visualized by Hoechst (blue) and Discs large (Dlg) is shown in magenta. Scale bars: 50 μm (A, C, E, G, I), 10 μm (B, D, F, H, J).

**Figure S5.**
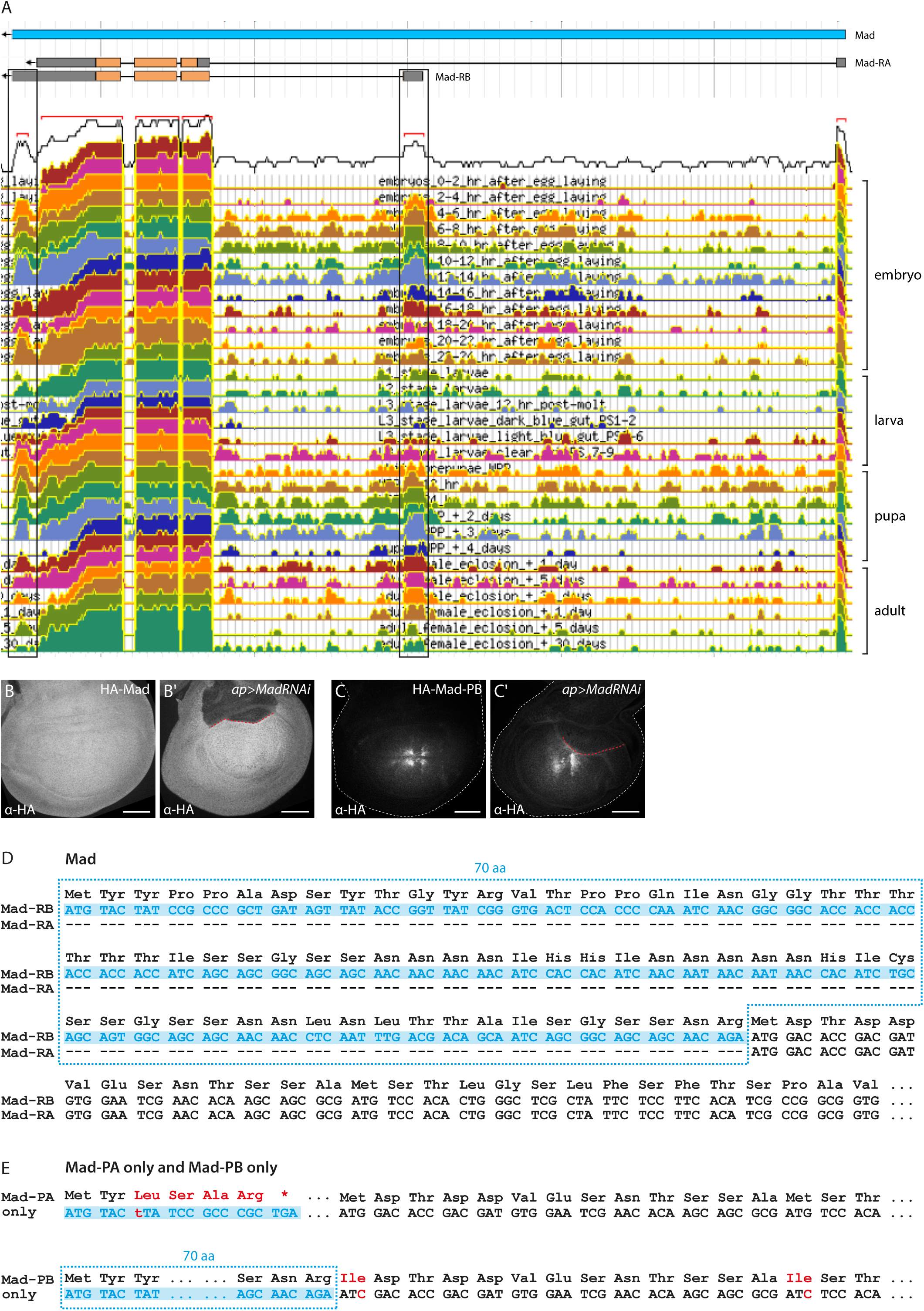
*Mad* isoform analysis. (A) RNA-Seq expression data for different developmental stages by the ModENCODE project predicts two transcript isoforms, Mad-RA and Mad-RB (Graveley et al., 2011). Screenshot from Flybase JBrowse and adapted. Black boxes highlight Mad-RB specific reads, labels at right indicate developmental stage. (B, C) Anti-HA staining of endogenously tagged Mad versions in late 3^rd^ instar wing discs. (B’, C’) RNAi-mediated depletion of Mad in the dorsal wing disc compartment using ap-Gal4 verifies specificity of the observed staining. Red dashed line indicates dorso-ventral compartment boundary. Scale bars: 50 μm. (D) Start of DNA and amino acid sequences of the two Mad isoforms. Mad-PB is an in-frame N-terminal extension of Mad-PA, containing 70 amino acids (marked in blue) upstream of the Mad-RA start codon. aa = amino acids. (E) To generate flies which exclusively contain one of the isoforms, we reintroduced mutated versions of the genomic sequence of *Mad* into *mad^[attP, KO]^* flies. For Mad-PA only flies, we introduced a frameshift into Mad-RB by inserting a single nucleotide. For Mad-PB only flies, we mutated the Mad-RA start codon (along with a second nearby ATG) by exchanging them with ATC (isoleucine). Changes compared to wildtype sequence are highlighted in red.

**Figure S6.**
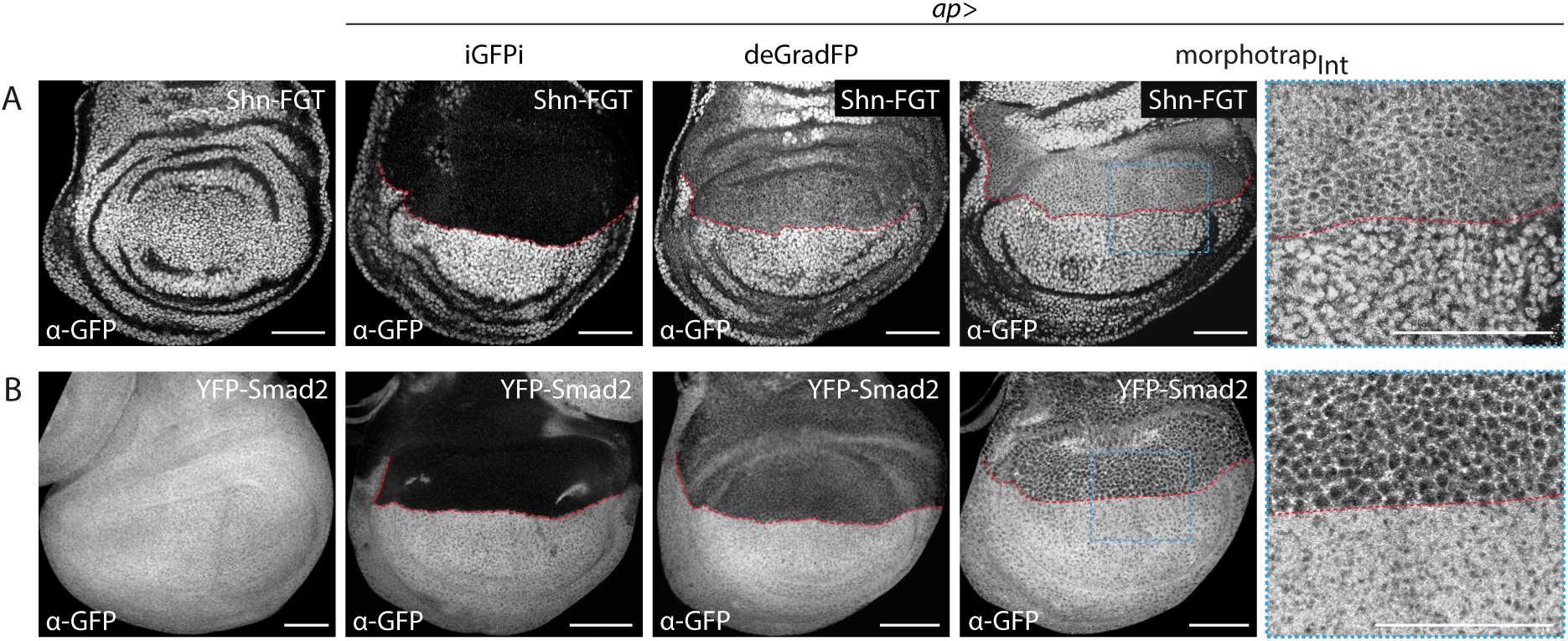
Tagged Shn and Smad2 can be manipulated with GFP-based tools. (A, B) Shn-FGT (A) and YFP-Smad2 (B) visualized by anti-GFP staining in wing discs either expressing the respective tagged allele alone or in combination with the indicated tool in the dorsal compartment using ap-Gal4. Red dashed lines mark dorso-ventral compartment boundary. Scale bars: 50 μm.

**Table S1:**
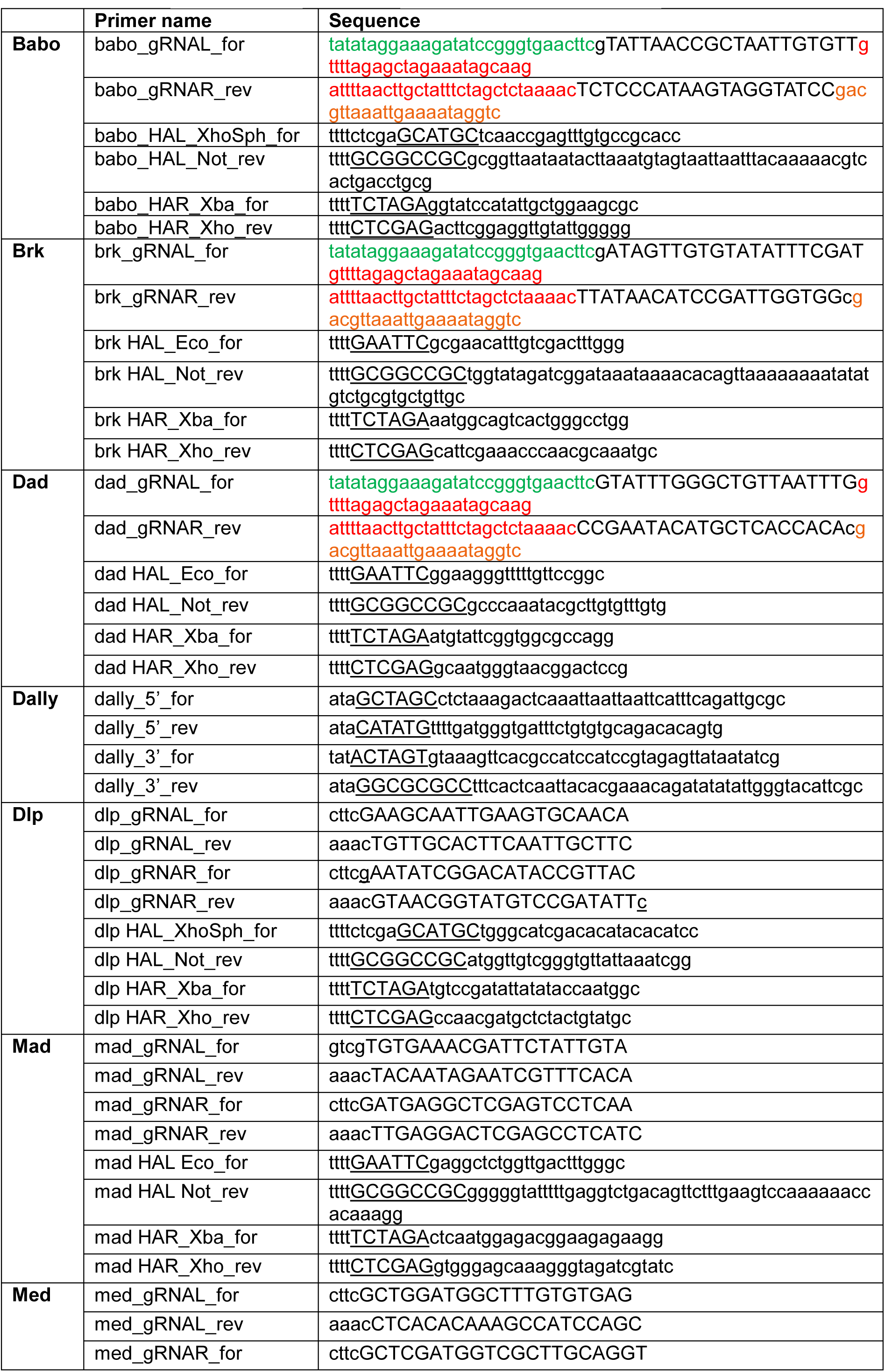
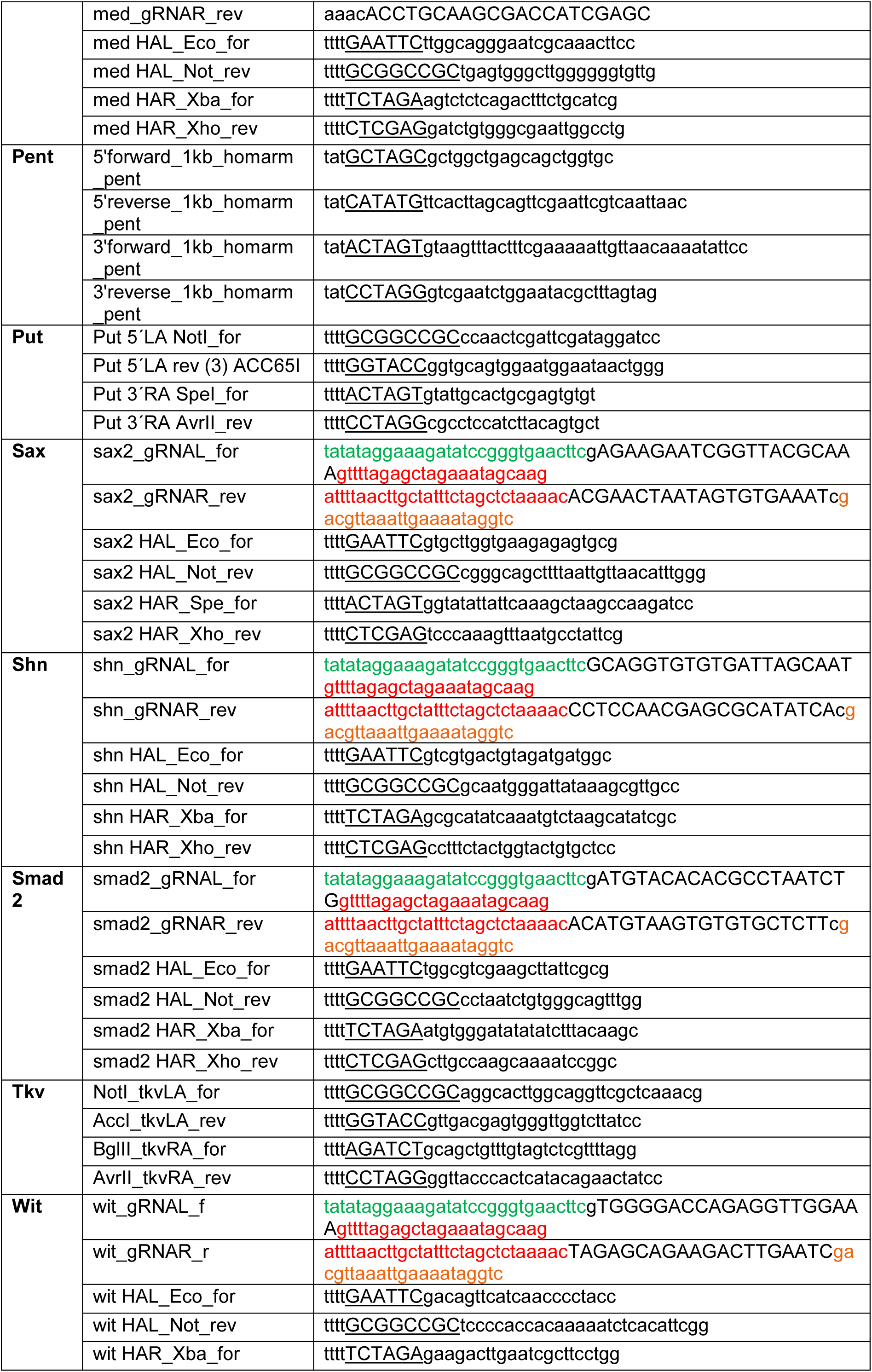
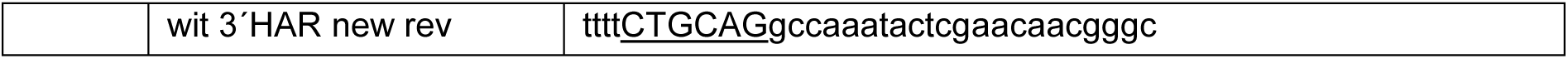
Primers to generate plasmids containing homology arms and guide RNAs.

**Table S2:**
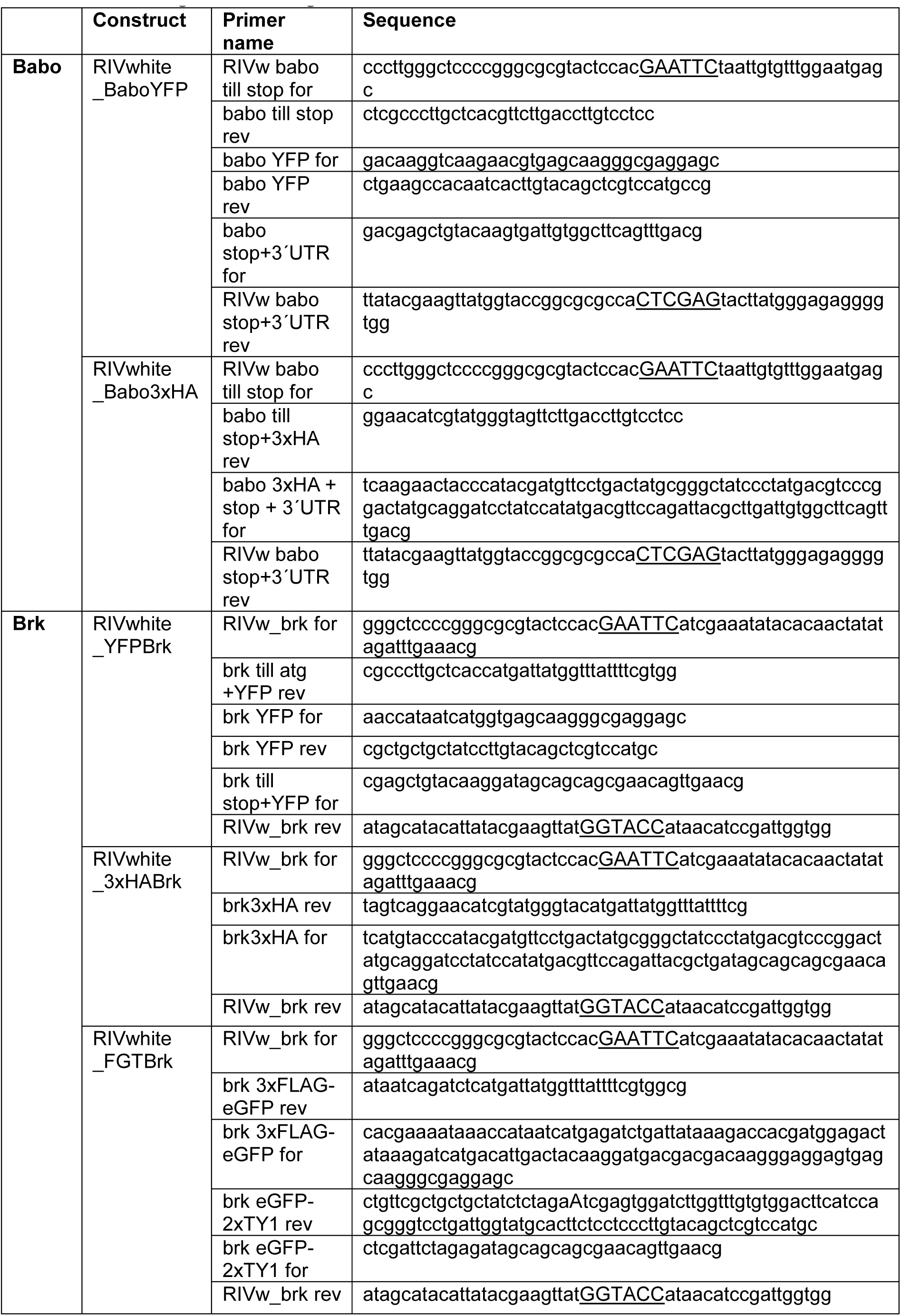
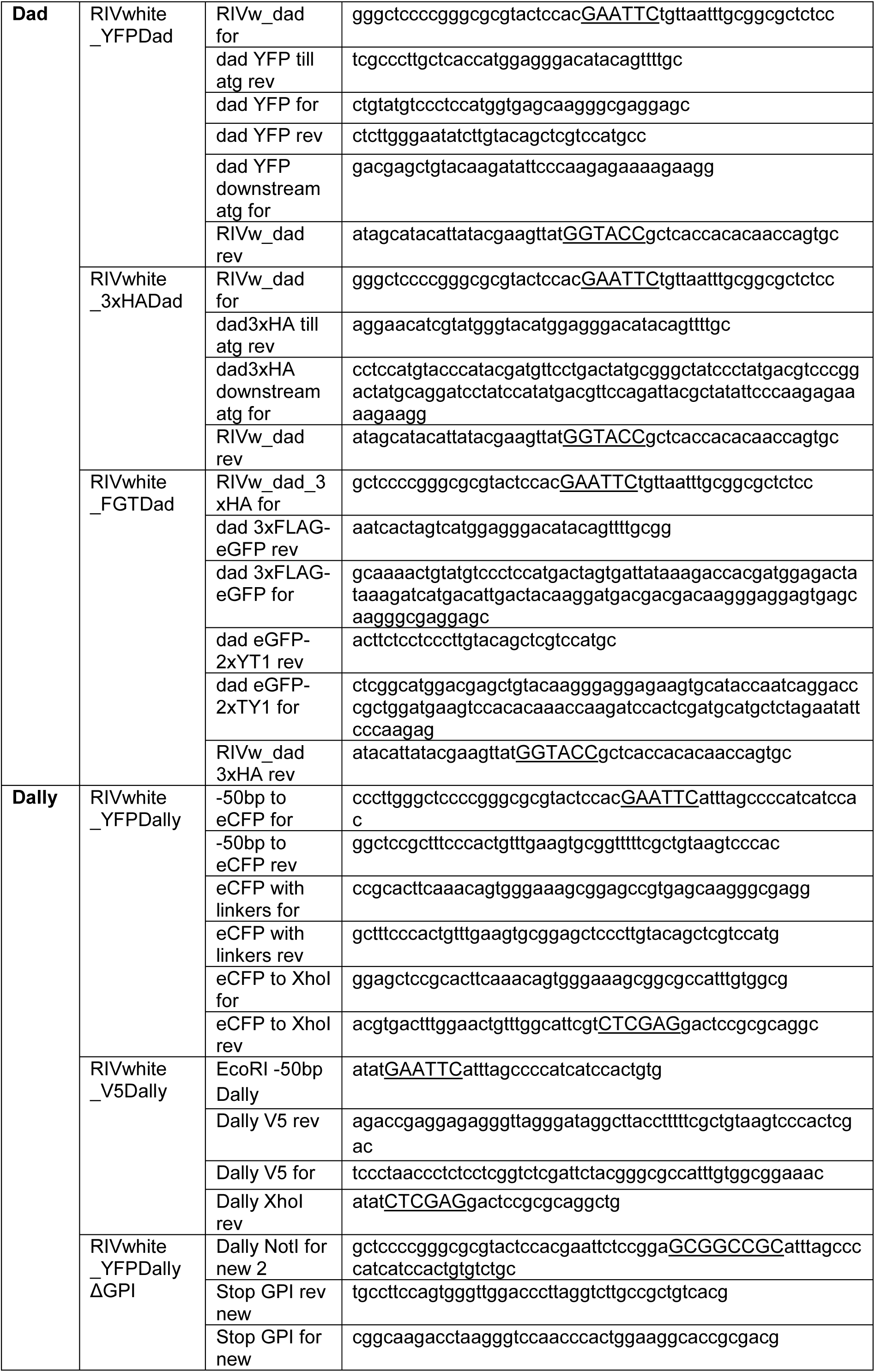
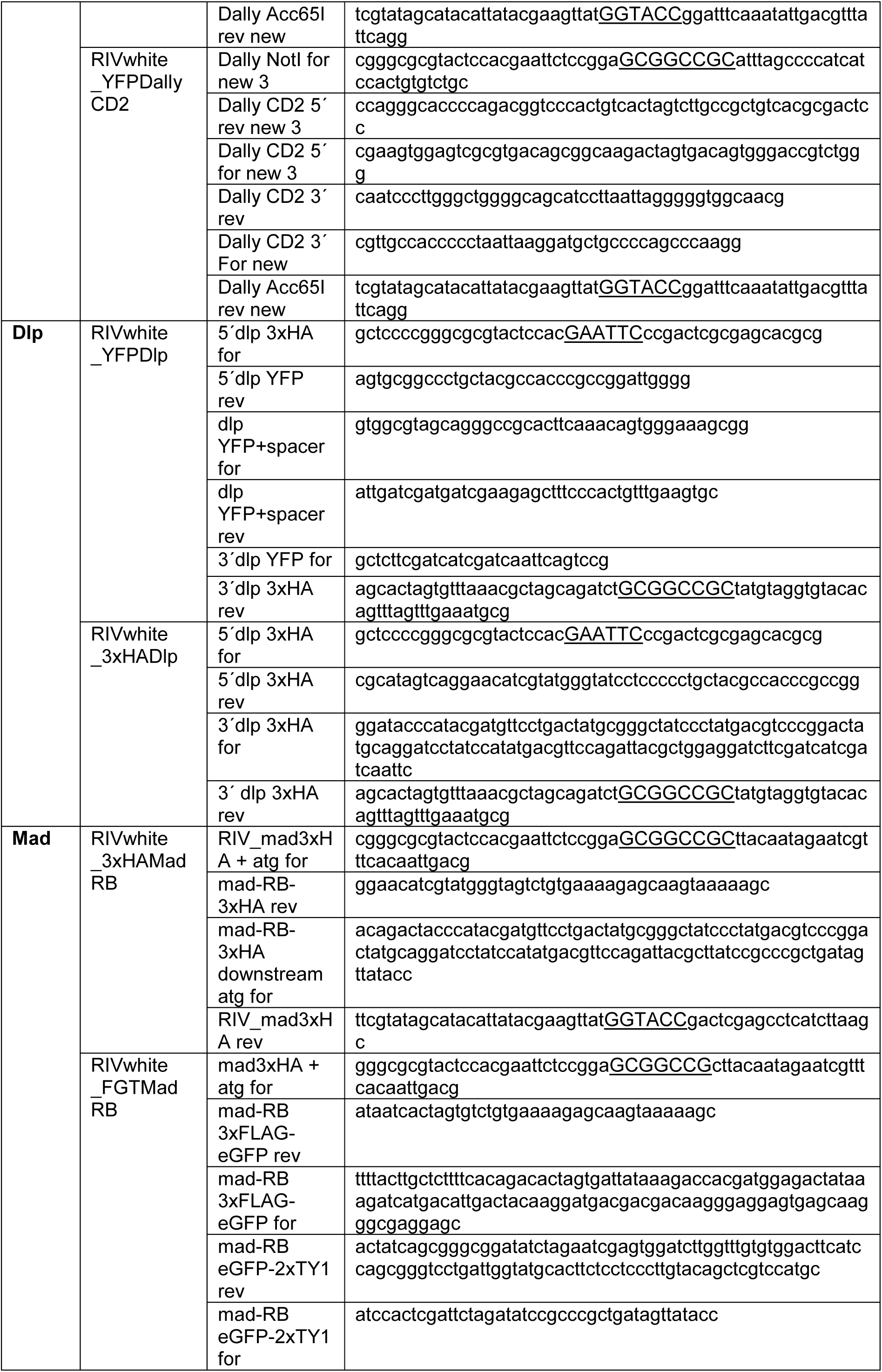
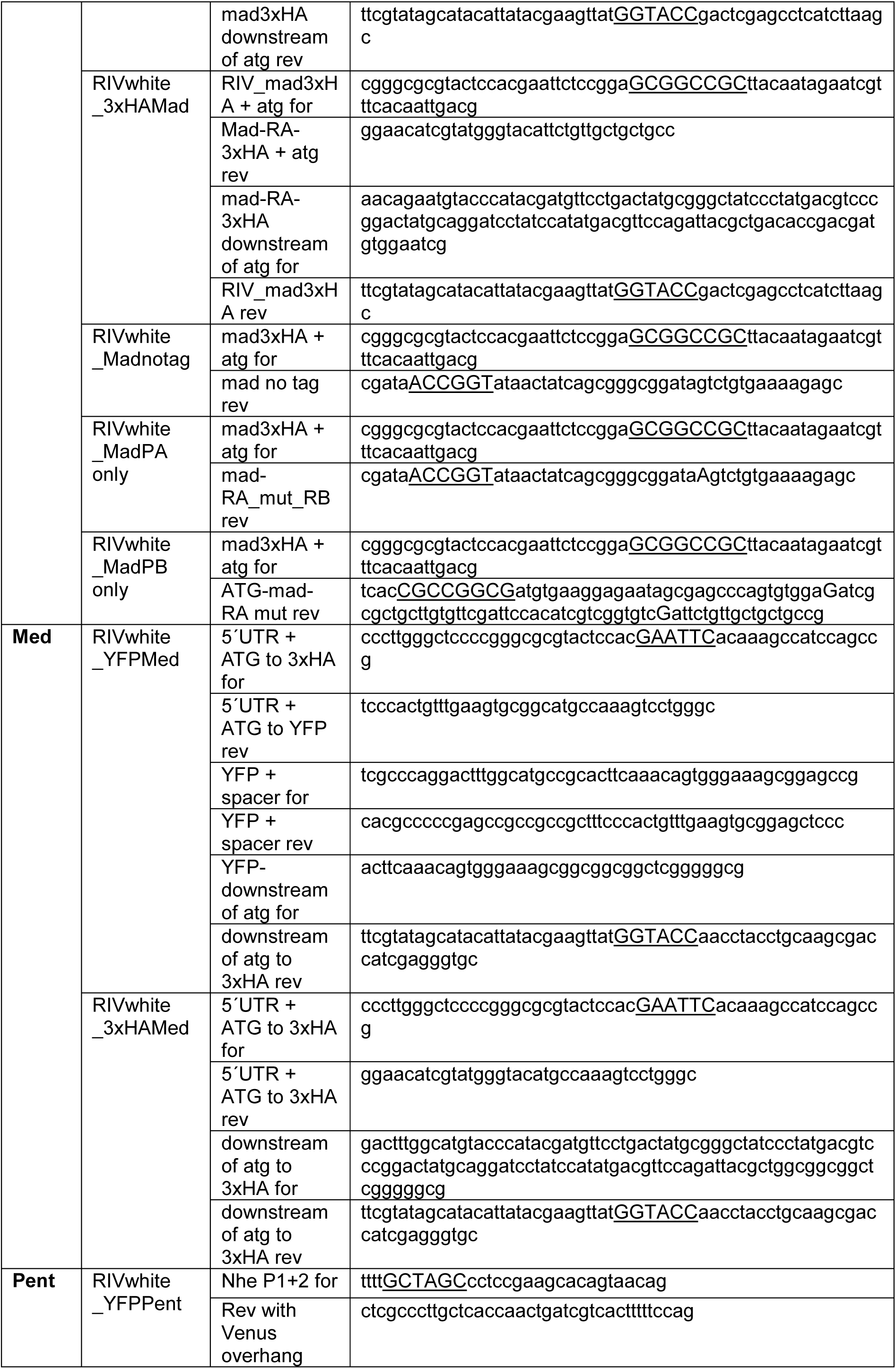
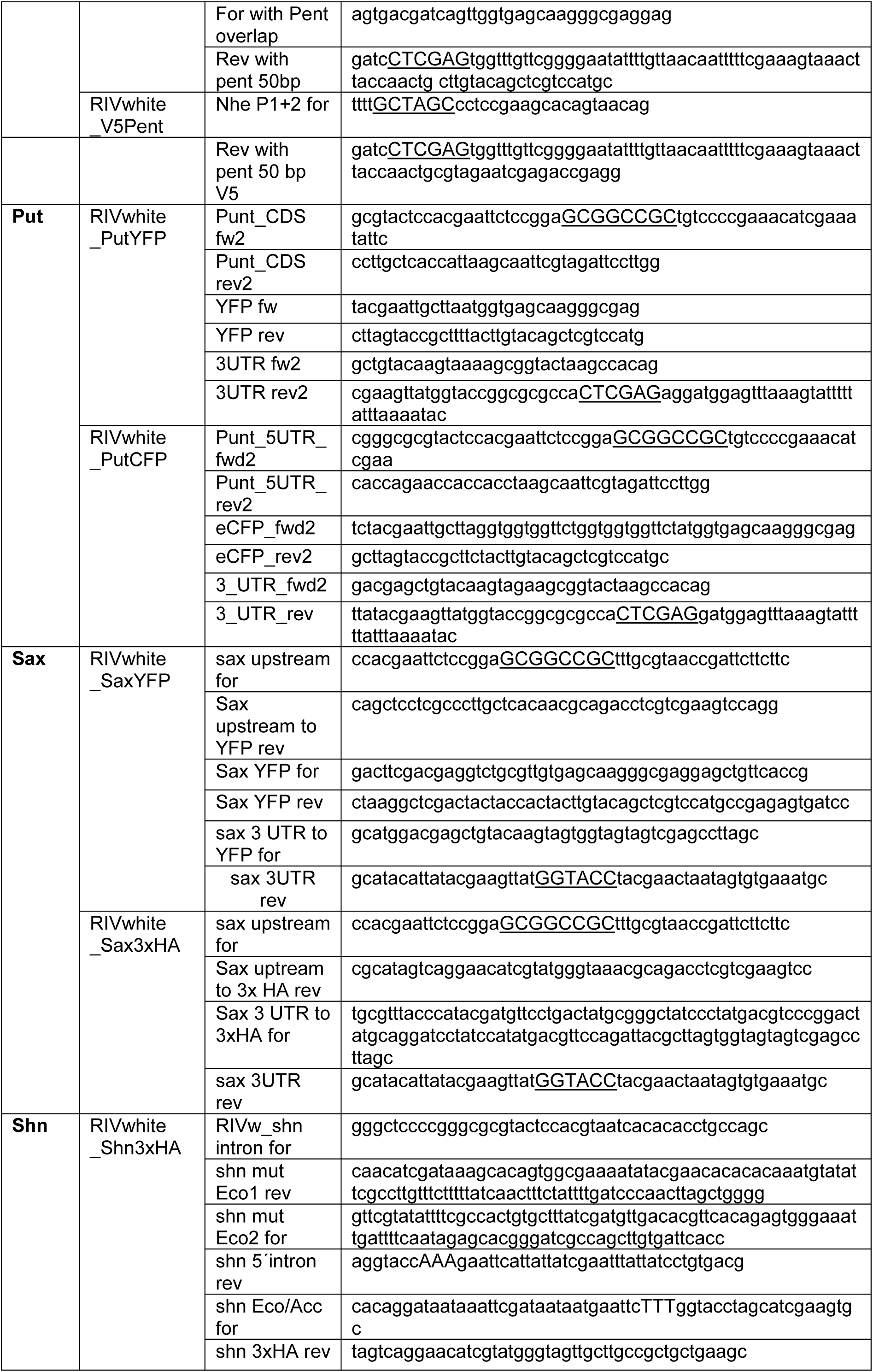
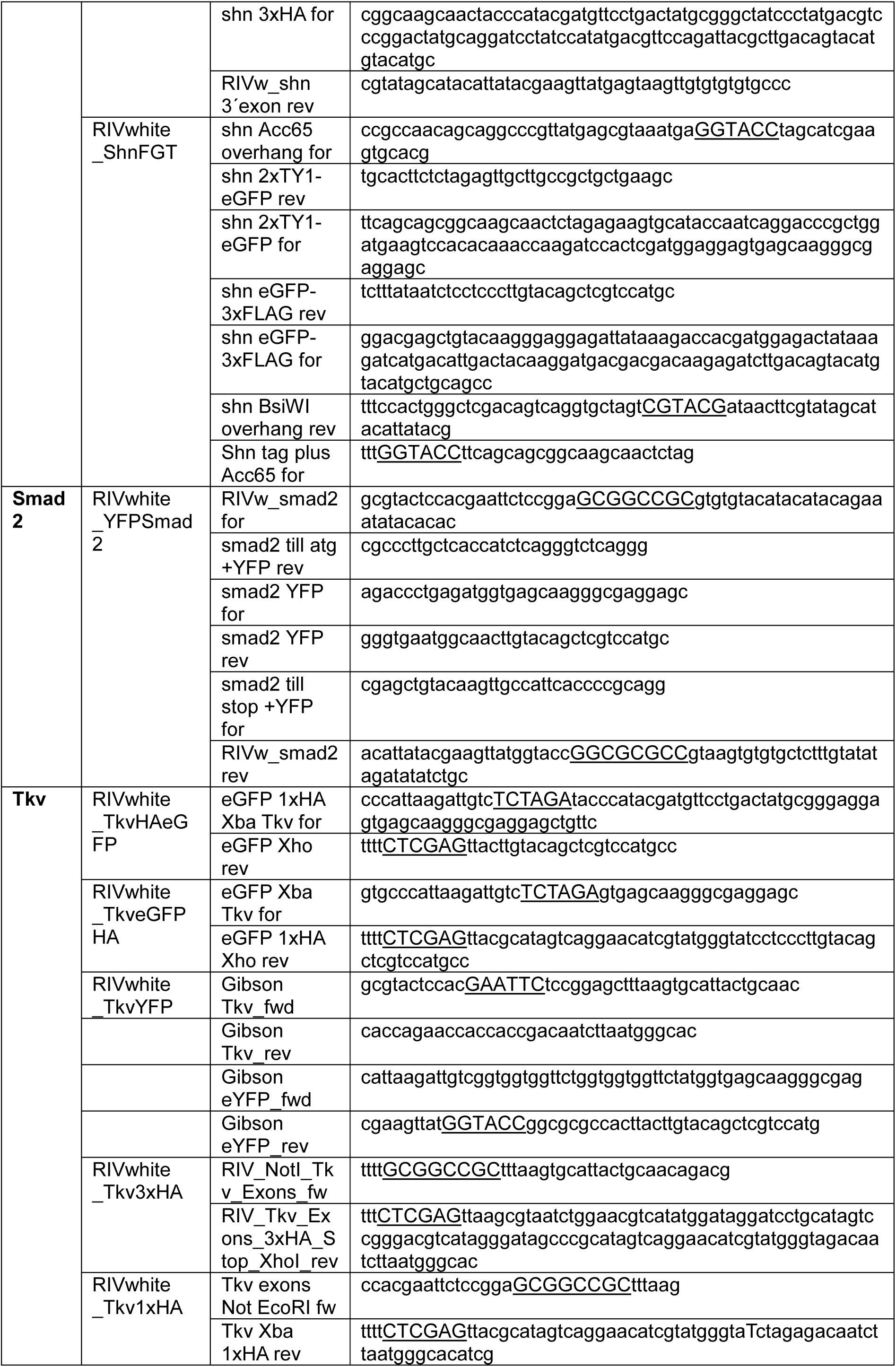
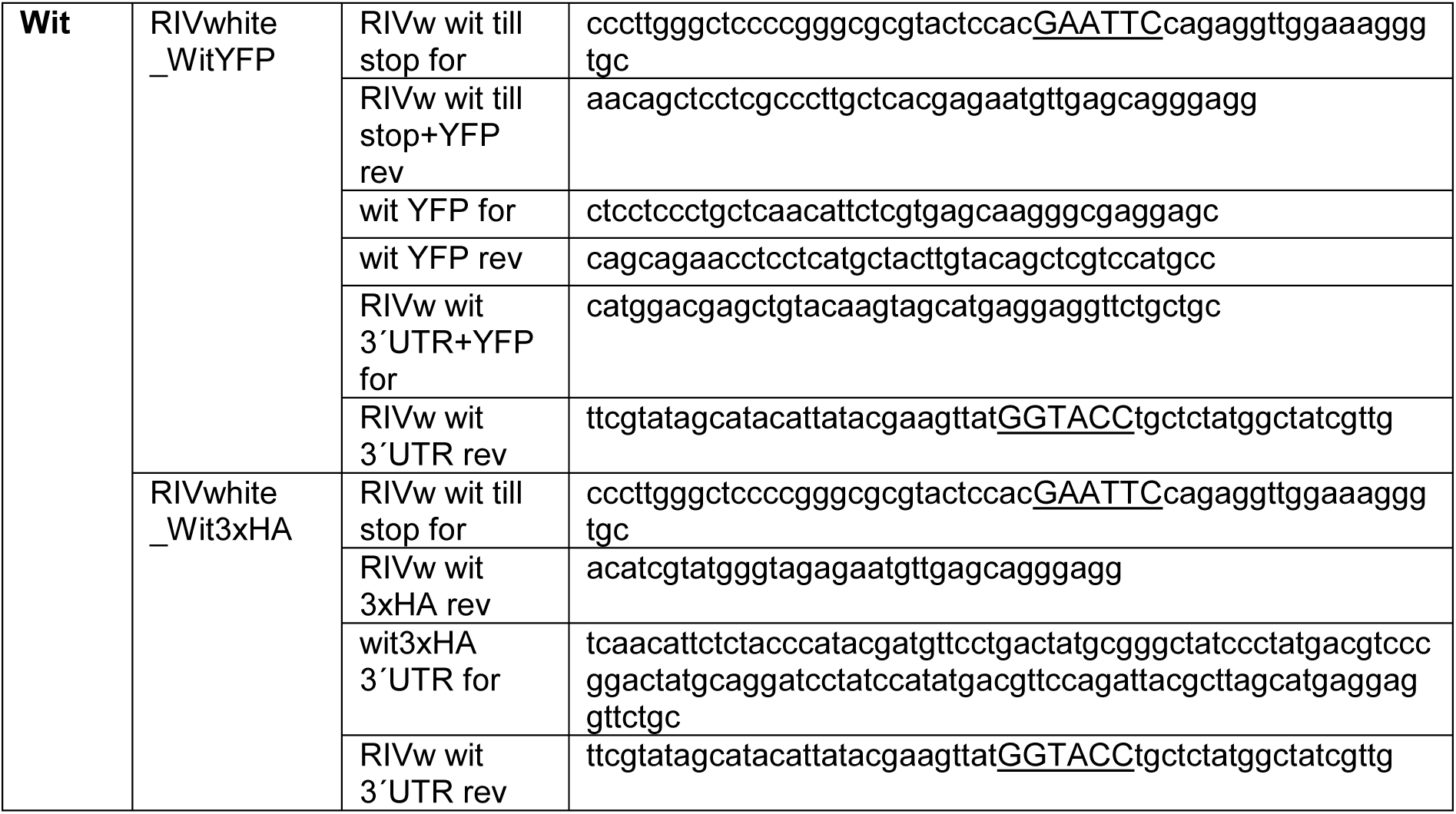
Primers to generate reintegration vectors.

**Table S3:**
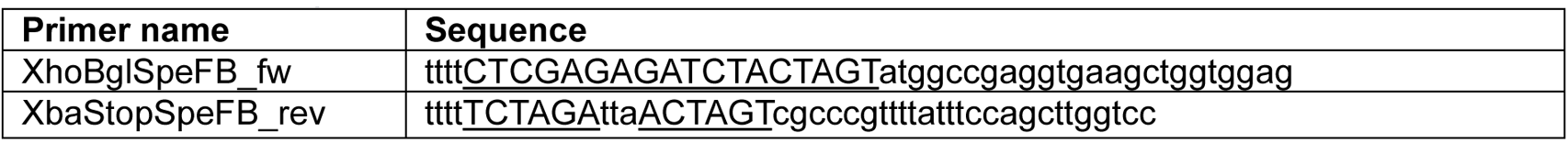
Primers to generate HA toolbox plasmids.

## Notes

### Competing Interest Statement

The authors have declared no competing interest.

